# Bridging geometric morphometrics to medical anatomy: An example from an experimental study of the human smile

**DOI:** 10.1101/2024.03.07.583999

**Authors:** Fred L. Bookstein, John T. Kent, Balvinder Khambay, Kanti V. Mardia

## Abstract

The method of Cartesian transformations introduced by D’Arcy Thompson a century ago in his celebrated book *On Growth and Form* precipitated an important development in 20th-century biometrics: a fusion of the geometrical and biological approaches to morphology. Some decades later this fusion, in turn, spun off another multidisciplinary focus, *statistical shape analysis,* that bridges between biostatistics and biomedical imaging. Our article is intended to seed a complementary focus: a bridge between biostatistics and medical anatomy, a field that has to this day remained mainly verbal rather than quantitative. Specifically, we are proposing a novel methodology for arriving at anatomical interpretations of statistical findings about large-scale contrasts of organismal morphology or its dynamics by combining two toolkits hitherto separate in their notation and their disciplinary housing: morphometrics and psychometrics. A contemporary morphometric analysis deals with patterns of shape coordinate covariation in terms of their geometric adjacency; psychometric factor analysis, the same patterns of covariation in terms of simplicity of interpretation. By combining these tools we account for the dynamics of a facial expression in terms of the actions of the underlying muscles, thereby realizing Thompson’s original metaphor, the “origins of form in force,” for systems that “vary in a more or less uniform manner.”

This paper reviews the history of Thompson’s metaphor and then the current literature quantifying smiles, in order to set the stage for the combination of scaling analysis and factor analysis that we are putting forward. We demonstrate the new approach by reanalyzing a data set of ten landmarks around the vermilion borders of the human lip contrasting the dynamics of two socially stereotyped physiological cycles, the open-lip smile and the closed-lip smile, over a sample of 14 normal faces. Our analysis centers on just two dimensions of statistical shape space, those of largest geometrical scale, which can be identified with the action of two different muscles, orbicularis oris and zygomaticus major. The two smiles differ radically in their achieved deformations. Furthermore, while the closed-lip smiles arrived at their final forms along similar shape trajectories, the open-lip smiles did not. Our closing discussion explores some aspects of morphodynamics that are illuminated by the example here.

---

> Epigraph:

> Shape does not admit of a definition in the language of real numbers.

> — Peter B. Medawar, 1945, page 157

## I. Introduction

Corresponding to our intention to build a bridge between two hitherto mostly separate disciplines, morphometrics and medical anatomy, this Introduction likewise divides into two parts, one for each.

### I.1. From D’Arcy Thompson to contemporary morphometrics

#### Transformation grids: D’Arcy Thompson’s version

D’Arcy Thompson’s masterwork *On Growth and Form (OGF)* of 1917 is, in Peter Medawar’s words, “beyond comparison the finest work of literature in all the annals of science that have been recorded in the English tongue. There is a combination here of elegance of style with perfect, absolutely unfailing clarity, that has never to my knowledge been surpassed” (Medawar 1958:232). There is no need, then, for us to begin by summarizing Thompson’s approach in Chapter XVII, “On the Theory of Transformations, or the Comparison of Related Forms”; better simply to use his own words.

Indeed Thompson is exquisitely clear about the thesis he wishes to establish: that a good grid diagram will permit, as a “comparatively easy task,” an assessment of “the direction and magnitude of the force capable of effecting the required transformation” (Thompson 1961:272). In the context of his examples, which are all comparisons across taxa, this goal became obsolete during the genomic revolution. But another thrust, that of dynamical simplification, is still cogent. Thompson sets two conditions: “that the form of the entire structure under investigation should be found to vary in a more or less uniform manner, after the fashion of an approximately homogeneous and isotropic body, … and that our structure vary in its entirety, or at least that ‘independent variants’ should be relatively few” (ibid., 274). From this he draws a sweeping conclusion:

> When the morphologist compares one animal with another, point by point or character by character, these are too often the mere outcome of artificial dissection and analysis. Rather is the living body one integral and indivisible whole, in which we cannot find, when we come to look for it, any strict dividing line even between the head and the body, the muscle and the tendon, the sinew and the bone. Characters which have differentiated insist on integrating themselves again; and aspects of the organism are seen to be conjoined which only our mental analysis had put asunder. The co-ordinate diagram throws into relief the integral solidarity of the organism, and enables us to see how simple a certain kind of *correlation* is which had been apt to seem a subtle and a complex thing.

> But if, on the other hand, diverse and dissimilar fishes can be referred as a whole to identical functions of very different co-ordinate systems, this fact will of itself constitute a proof that variation had proceeded on definite and orderly lines, that a comprehensive ‘law of growth’ has pervaded the whole structure in its integrity, and that some more or less simple and recognisable system of forces has been in control. It will not only show how real and deep-seated is the phenomenon of ‘correlation,’ in regard to form, but it will also demonstrate the fact that a correlation which had seemed too complex for analysis or comprehension is, in many cases, capable of very simple graphical expression. (ibid., 275–6; emphasis Thompson’s)

The key idea, then, was to work with pairs of drawings of organisms rather than with measurements. To show the comparison, Thompson would draw a grid of vertical and horizontal lines over the picture of one organism in some orientation, then deform these verticals and horizontals over the picture of a second organism so that grid blocks that corresponded graphically would roughly correspond morphologically as well. He liked to interpret these graphical patterns via a metaphor, the explanation of form-change as the result of a “system of forces.”

Perhaps the most familiar example of Thompson’s method is his comparison of the porcupine fish *Diodon* to the sunfish *Orthagoriscus mola.* His drawing is reproduced in our Figure 1. Here, Thompson explained,

> I have deformed [the *Diodon* grid’s] vertical coordinates into a system of concentric circles, and its horizontal coordinates into a system of curves which, approximately and provisionally, are made to resemble a system of hyperbolas. The old outline, transferred in its integrity to the new network, appears as a manifest representation of the closely allied, but very different looking, sunfish *Orthagoriscus mola.* [This] particularly instructive case … accounts, by one single integral transformation, for all the apparently separate and distinct external differences between the two fishes. (Thompson 1961:300)

**Figure 1.**
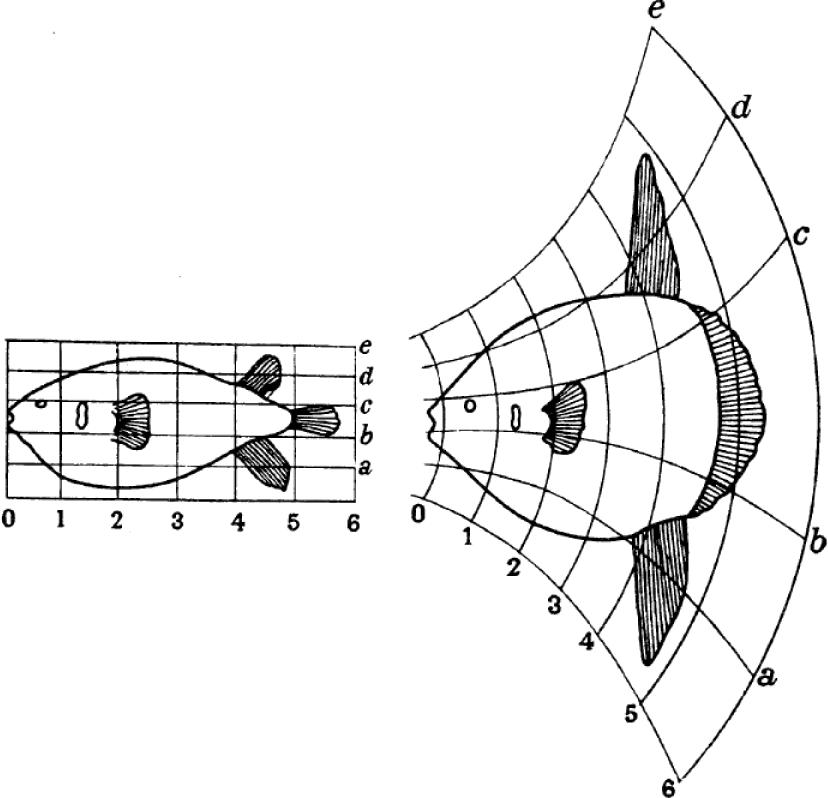
The most famous of Thompson’s examples: *Diodon* (porcupine fish) to *Orthagoriscus* (sunfish). After Thompson (1961:301).

One of us (Bookstein, 1978) has reviewed the “vicissitudes” of this insight of Thompson’s from its inception in 1917 to the dawn of contemporary geometric morphometrics (GMM). It is generally conceded that, again in Medawar’s words,

> The reason why D’Arcy’s method has been so little used in practice (only I and one or two others have tried to develop it at all) is because it is analytically unwieldy. (Medawar 1958:231)

Thompson’s examples often involve ludicrously distorted forms (e.g., the “human skull” on page 318 of the 1961 edition) or systematically misrepresent the matching of identifiable locations between the grids of a pair, as in Figure 1. The grid on the right does not “account for” those “external differences” in any quantitatively useful way (Bookstein 1978:70–71, 124–128) — it is just not accurate enough. Nor does a more exact rendering aid the situation, as when the diagrams are more carefully produced, “not only are the grid lines deformed in several ways, but the deformation is different in different parts” (Sokal and Sneath, 1963:83). In other words, when these grids are accurate, they cannot be reduced to words; when they are legible, it is because they are inaccurate.

#### Peter Medawar’s version

But the critique we have in mind for this essay focuses on a deeper issue than was treated in Bookstein, 1977, 1978. Can a closer study of the terms like “simple” and “integral” that drive Thompson’s metaphor suggest tools for dissecting separably explicable components of the “system of forces” that is ostensibly to have “been in control”? Back in the 1940’s Medawar had already sensed the strange reversal of description and explanation entailed in Thompson’s suggestion:

> The purpose of [Thompson’s] analysis is primarily and fundamentally descriptive. The need for an accurate technique of descriptive analysis is justified by the belief that a knowledge of *what happens* in any physiological process should precede an inquiry into how it comes about. (Medawar 1944:139; emphasis Medawar’s)

This “knowledge of what happens” turned out to require an extension of the language of quantitative morphological description, *and that extension requires an acknowledgement of the explanations that will ultimately follow from the graphics*.

In speculating on improvements to Thompson’s method, Medawar touched on the two main techniques that drive the contemporary synthesis to be sketched in the next few pages. One enduring theme, the separation of considerations of size change from those of shape change, was originally set down in a letter he wrote to Thompson in 1942: that the role of geometry should be consistent with Felix Klein’s approach (the “Erlangen programme”) via invariant properties under groups of transformations. Medawar (1944) explicitly embraced this principle in his own data analysis of human growth when he scaled each growth stage to the same total height on the printed page. In proceeding so, Medawar just missed the idea of “shape space” for the same data, as would have been represented by the one-dimensional formalism of Small (1996:14–17) half a century later. Note, too, how Medawar switched tacitly from evolutionary to developmental applications likelier to lead to simpler reports of findings.

But there was a second pole of this evolution that Medawar got completely wrong: the relevance of the two- or three-dimensionality of the world that we and the organisms we study all live in. Medawar analyzed the longitudinal dimension of human growth, and then commented, “The dimension of height will here be called *x* and of breadth *y*; they may be dealt with separately” (Medawar 1944:135). In that assertion he was badly mistaken. Huxley (1932), for instance, had already noticed that even if the sides of a rectangle obey his allometric law (with which Medawar did not agree), its diagonal need not — the analysis of a shape change requires that substantial attention be paid to the ways that deformations of square Cartesian grid *cells* can actually vary, not just what the individual grid *lines* might look like (“circles” and “hyperbolas,” in Thompson’s *Diodon* example). Descriptions of this type have proved scientifically worthless; we need a principled algorithmic substitute.

#### The emergence of geometric morphometrics

The same 1978 essay by Bookstein that sharpened the critique of *OGF* went on to suggest a replacement technique, the biorthogonal grid, that, in the event, did not conduce to a sound biometrical toolkit. (In the modern context it applies only to the uniform term of the transformation.) The key proved instead to be a mathematization of the *rhetoric* that Thompson was intuiting, rephrased for a context of signal processing that explicitly accommodates ideas of both true biological variance (which, remember, Thompson explicitly refused to study) and noise (as of landmark location). The method of transformation grids thereby turned into the praxis today universally known as *geometric morphometrics* (GMM), which has three core components:

i. a space of shapes of landmark point configurations as equivalence classes (Kendall, 1984), together with two metrics, one (Procrustes distance) geometric and a-priori, the other (covariance structure) biometric and data-based;
ii. a pattern engine for coordinate transformations (the thin-plate spline, which automates the drawing of the transformed coordinate grid on which the interpretation of factors depends); and
iii. a protocol for extracting component grids, together summing to an observed transformation, that individually admit potentially valid biological explanations: machinery for getting from arithmetic to understanding by multiple projections.

The GMM method is evidently designed for data in the form of discrete landmark locations. Our eyes and our machines also attend to curves, surfaces, textures, and a variety of other visual resources. Methodologies for those resources lag behind the easy ability of the landmark approach to embrace the methods of covariance analysis that drive applications in other domains of moderate complexity, such as econometrics or psychometrics. For state-of-the-art tools for curves, see Srivastava and Klassen, 2016, and, for gray-scale images, Grenander and Miller, 2007; but this tradition emphasizes classification or detection of anomalies, not the dissection of collections of empirical examples with an eye toward rigorous assertions about biological causes.

It must be acknowledged, then, that this methodology, while responding explicitly to Thompson’s challenge, does not parallel Thompson’s actual *method* in any coherent way. Thompson was concerned with outline drawings, GMM with landmark configurations, which are a far more stylized measurement channel; Thompson attended to the shape and spacing of the coordinate curves per se, GMM to the sizes and shapes of the little grid cells as they deviate from the forms of their neighbors; Thompson was interested in pairs of forms, whereas GMM embraces analysis of patterns across samples of arbitrarily many landmark configurations. Most saliently, to Thompson there is only one Cartesian transformation per biological comparison, and it *is* the explanation; whereas in GMM, transformations are considered in groups, as sets of potentially helpful diagrams that might lead the investigator to explanations later that take the form of group comparisons, trends, growth-gradients (or their complement, the focal transformations), responses to variations of the environment, or any other coherently reported cause. What our grids have in common with Thompson’s is this powerful cognitive property of *suggesting* explanations. The price of preserving that power was the transfer of the technique out of Thompson’s näıve setting, the single organism-to-organism comparison, into the rigorous context of statistical signal processing, a praxis that did not even exist until Thompson’s extreme old age. Several monographs survey this GMM toolkit, for instance, Dryden and Mardia, 1998, 2016 and Bookstein, 2014, 2018, and there are also many workshops and short courses for the interested biologist to choose from.

#### GMM’s standard dataflow: shape coordinates

The setup for the standard GMM portion of the dataflow for today’s example is set out in Figure 2 and the right-hand column of Figure 4. The data resource driving all the arithmetic is the set of video coordinates of landmark decagons (configurations of ten detached points) in two conditions from one of our 14 experimental subjects as referred to a synthetic frontal view (Figure 2, left column; their anatomical origin will be reviewed later). The upper row of diagrams corresponds to the first experimental condition; the lower row, the second. There is an overall vertical shift of these points because the action of smiling shifted their center of gravity with respect to the solid head. In the central column the data have been centered, decagon by decagon, thus correcting for this specific translation in the laboratory coordinate system, which is of no interest here. Finally, the right-hand column shows the conversion to *Procrustes shape coordinates,* which standardize the decagons of the center column to a least-squares superposition on their average with Centroid Size (sum of squared distances from the centroid) 1.0 and principal moments horizontal and vertical. Whichever column we are examining, the two smiles look different; our task is to provide an anatomical explanation for this subject (and, it will turn out, twelve of the thirteen others).

**Figure 2.**
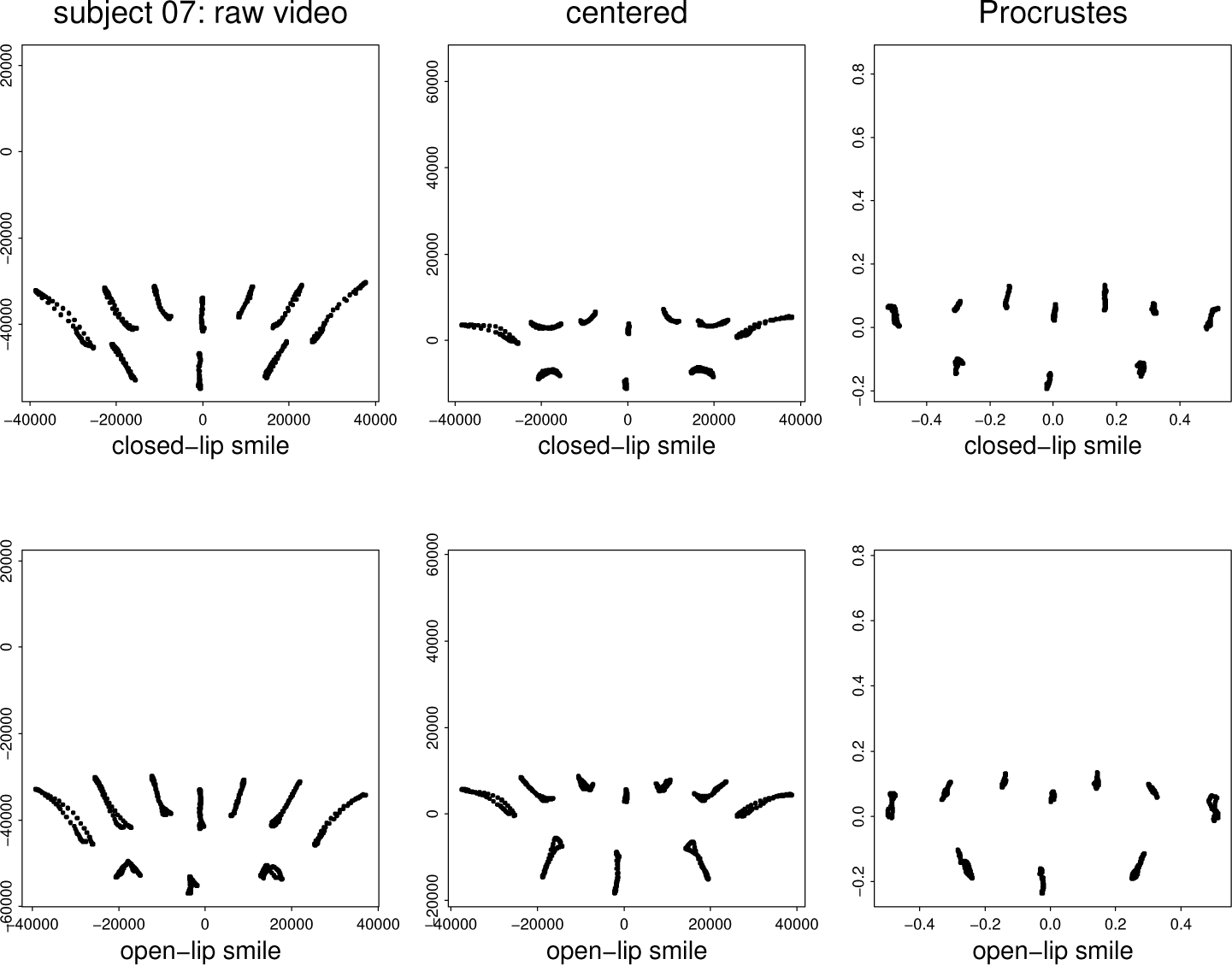
The standard first steps in a GMM analysis, as applied to the data from one of our little study’s fourteen subjects. See text. These and all other statistical computations in this paper went forward in the standard statistical package Splus, a dialect of R.

#### From shape coordinates to partial warps

Depending on the biomathematical question at hand, any of the superpositions in Figure 2 might have been appropriate (it will turn out to be an extension of the right-hand column that we will be using). To choose among such a series of standardizations is the crucial step in the contemporary rationalization of Thompson’s method. (Thus Medawar’s handwritten note to Thompson in 1942 anticipated GMM by 50 years.) What are rendered here are *equivalence classes* of forms, not forms per se. Throughout GMM that is how our biometric statistical analyses now universally go forward. In the extant literature applying GMM to smiles, as reviewed in Section I.2, the biomathematics is limited to that of Figure 2 only: means and variances of the coordinates of the landmarks in the right-hand column.

To bridge to the science of medical anatomy, however, we need (forgive the metaphor) a sharper scalpel. We will be emphasizing one of the more sophisticated (thus less often invoked) tools from today’s GMM toolkit, the decomposition of any observed Cartesian transformation into its *partial warps*. Any thin-plate spline deformation of one landmark configuration onto another has a *bending energy* that is computed by preand postmultiplying a *bending-energy matrix BE* first by a vector of the *x*-coordinates and then by a vector of the *y*-coordinates of the target configuration, and finally summing the two evaluations. (The matrix *BE* is the quadratic form representing the summed squared second derivatives of the corresponding thin-plate spline interpolation, integrated over the entire picture plane: see Bookstein, 1991.) This bending-energy matrix *BE* is a function only of the starting form for the deformation (maybe a textbook example, maybe your sample average). It is *p* × *p,* where *p* is just the count of landmarks in your data set. *BE* has *p* − 3 nonzero eigenvalues corresponding to the *p* − 3 geometrically orthogonal modes of “bending” of the configuration, that is, the *p* − 3 orthogonal ways that either just the *x*− coordinates or just the *y*−coordinates can be transformed so that a grid of squares does not transform into all the same parallelogram under the action of the thin-plate spline whose bending-energy matrix this is. The corresponding *p* − 3 eigenvectors are the *principal warps* of the transformation, and the projections of the sample shape change onto these eigenvectors, first in the *x*-direction and then in the *y*-direction, are called the *partial warp scores*; they make up one 2-vector for every specimen in your sample. To this list of *p* − 3 pairs of scores we usually add a “zeroth,” the *uniform component* of these same transformations, that describes its *affine* changes, those that shear squares into the same parallelogram all over the starting grid. (For its formula, see Dryden and Mardia, 2016, p. 290, or Bookstein, 2014, p. 414.) The total is thus *p* − 2 complex numbers, or 2(*p* − 2) scalar scores. These counts are all for the case of 2D data. For data in 3D, all those terms *p* − 3 change to *p* − 4, and the partial warp scores are now real 3-vectors, except for the uniform term, which now has five components.

For the two smiles in Figure 2, these decompositions are set out in Figure 3. Across the top are the 10 − 2 = 8 partial warps for the average configuration, each one realized as a thin-plate spline transformation grid corresponding to the appropriate projection of the deformation (treated as a 10-vector of complex numbers) tracking the form of the open-lips smile from its rest configuration to its configuration of maximum expression. (For the definitions of these terms, see Section II.) Each of these deformations has been exaggerated by a factor of two to enhance its legibility. Along the middle row are the scatterplots of the corresponding pairs of coefficients frame by frame along the trajectory of the open-lips smile. The leftmost column is for the uniform term, while columns 2 through 8 survey the first, second, …, seventh *nonuniform warps,* having specific bending energies 2.8, 9, 11, 23, 27, 48, 73. These numbers roughly correspond to a scale of spatial focus (Bookstein, 2015a). It is clear from the scatters that the signal amplitude of these terms is roughly comparable between the zeroth and first (columns 1 and 2) and then falls off drastically for the remaining six pairs of dimensions. For the closed-lips smile (lower row), the dominance of those first two columns of scatters persists, but the trajectory of the uniform component over the motion is different from that of the other smile, and likewise that of the fourth partial warp.

**Figure 3.**
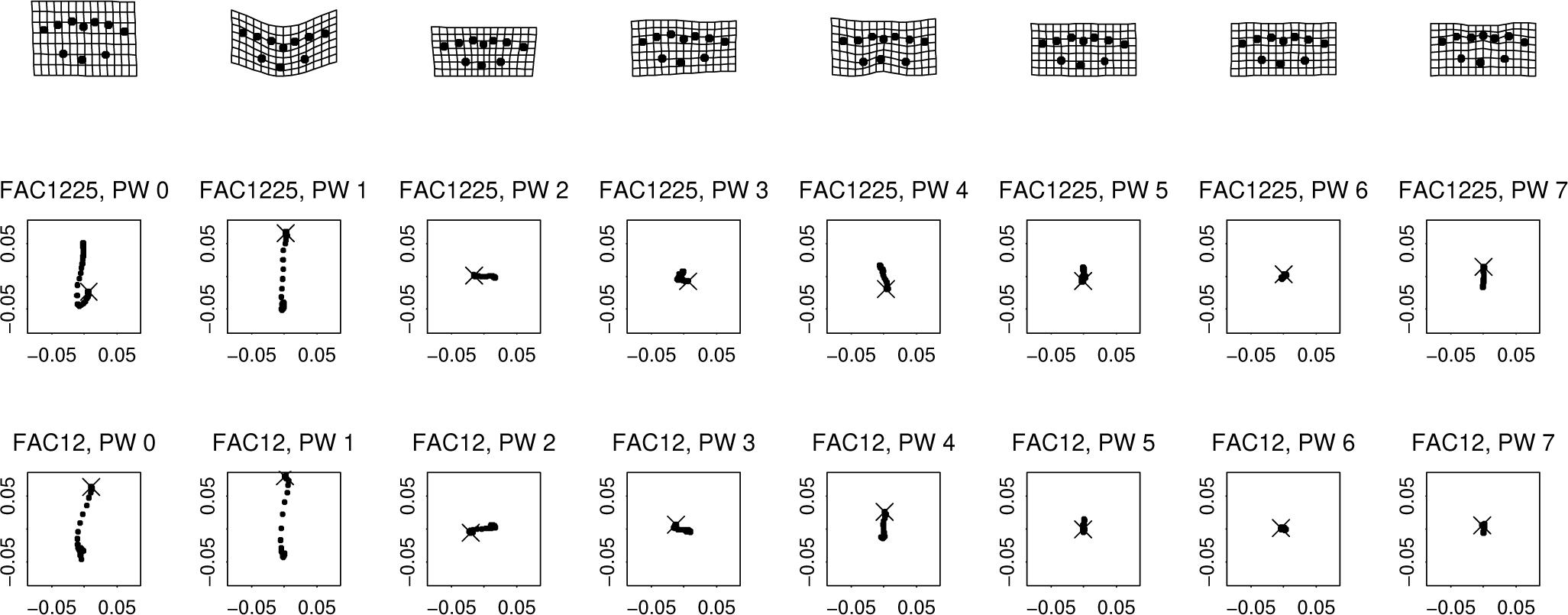
Partial warp analysis of the trajectories of decagonal shapes (Figure 2, lower right) between rest and the configurations of greatest smile expression for subject 07’s smiles. (top row) The eight components drawn as double the mean shift from rest position to the configuration of greatest open-lips smile expression. (middle row) The corresponding scatters. In each frame the rest (starting) position is indicated by an ×. Nearly all of the signal is on the first two of these components. (bottom row) Scatters for the closed-lip smile, same subject.

Both of these notions, the principal warps and the partial warp scores, must be distinguished from the *relative warp scores,* which in the present context are the same as the ordinary principal component scores of the sets of shape coordinates in figures like Figure 2 here, interpreted just as 2 × 10 = 20 ordinary variables. They are also the principal components of the partial warp scores themselves, the 16 Cartesian coordinates in Figure 3. Corresponding to these scores there is also a suite of 2 × 10 − 4 = 16 vectors of loadings, the *relative warp loadings*. The partial warp scores are just a rotation of the original Procrustes coordinates; relative warp analysis often goes better in terms of that basis (cf. Figures 13 and 14).

The relative warp analysis of this same subject’s smiles, Figure 4, conveys a surprise of which the biomathematical interpretation is the main subject of Section III. For the open-lips smile, and that smile only, the scatter of the zeroth and first partial warps is indistinguishable from a rotation of the first two standard GMM relative warps. By identifying the plane of these two relative warps with this dominant sublist of the spectrum of spatial scales, we take our first step along the bridge from all this GMM arithmetic toward a valid anatomical interpretation.

**Figure 4.**
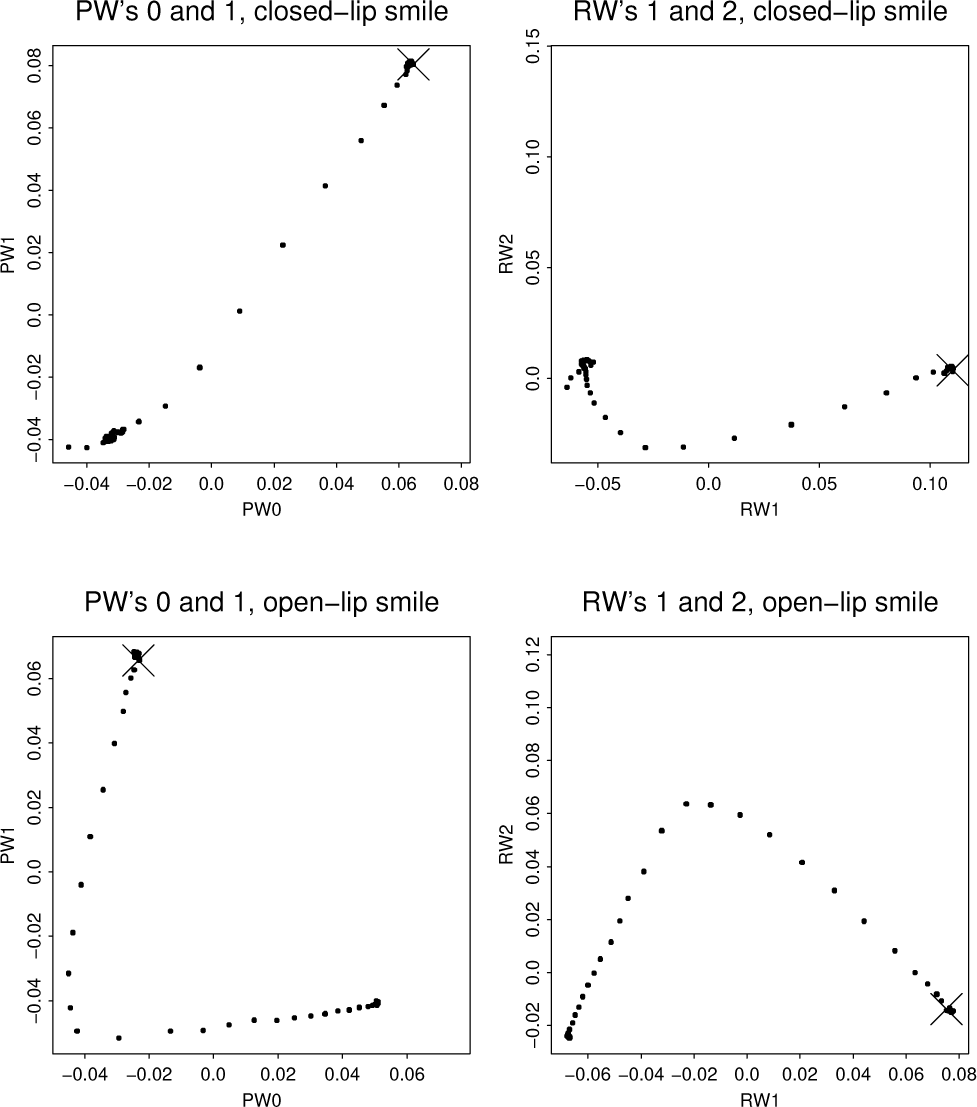
Still restricting our attention to this one single subject, we notice that the standard GMM relative warps analysis of either smile conveys the same information as just one end of the spatial spectrum of these features. But the smiles certainly differ in their configuration within that subspace. Section III is devoted to working through this paradox and its anatomical implications for the full data set of fourteen subjects. × symbols indicate rest (starting) position as in Figure 3.

I.2. Aperçu of quantitative anatomy: analysis of human smiles

In *OGF* the claim that changes of form originated in force was at best a metaphor. Thompson left it an open question whether there were examples of *actual* “simple and recognisable systems of forces” leading to *actual* changes of form along “definite and orderly lines.” A suitable testbed for exploring such a possibility would be a setting for which that “system of forces” was already known from the anatomy textbooks. To set the context, we briefly summarize the current literature regarding quantification of human smiles. (We do not pretend to review the topics of the *simulation* of smiles, as by avatars, or their recognition in images, as in industrial biometrics; those are larger literatures only tangentially related to our topic here.)

The human smile, one of the most fundamental modes of interpersonal communication, has been the subject of experimental study ever since Darwin first studied its empirical role in this regard (Jabr, 2010) in connection with his more general comparative explorations of this topic. Indeed Darwin’s great treatise on “expression of emotions” devotes many pages to the contemplation of smiles (e.g., perhaps “the habit of uttering loud reiterated sounds from a sense of pleasure first led to the retraction of the corners of the mouth …,” p. 208). Only later did the sciences of anatomy and physiology extend to the analysis of the modes by which this evocative facial expression is produced, specifically, the elementary muscle actions conveniently reviewed in Rubin, 1974. The topic can be categorized as one corner of the general three-dimensional study of facial soft tissues (see reviews in Bowman, 2008 or Sforza et al., 2013).

The biomathematician needs to begin with the sort of diagram exemplified here in Figure 5, showing the anatomical origins of all the relevant information in terms of a combination of Latin names and image properties. Plainly the human smile involves numerous muscles. One group, including zygomaticus major, moves the corners of the mouth upwards, outwards, and backwards, resulting in the “closed-lip” or “lips-together” smile. A second group of muscles that variously insert on orbicularis oris around its orbit contribute to a smile by separating the lips, producing a “full-lips-apart-tooth-showing smile,” which here will be called the “open-lip smile.” (A few sources refer to this as the “maximal smile” instead.) Besides the zygomaticus major these other muscles include levator labii, depressor labii, risorius, zygomaticus minor, and others.

**Figure 5.**
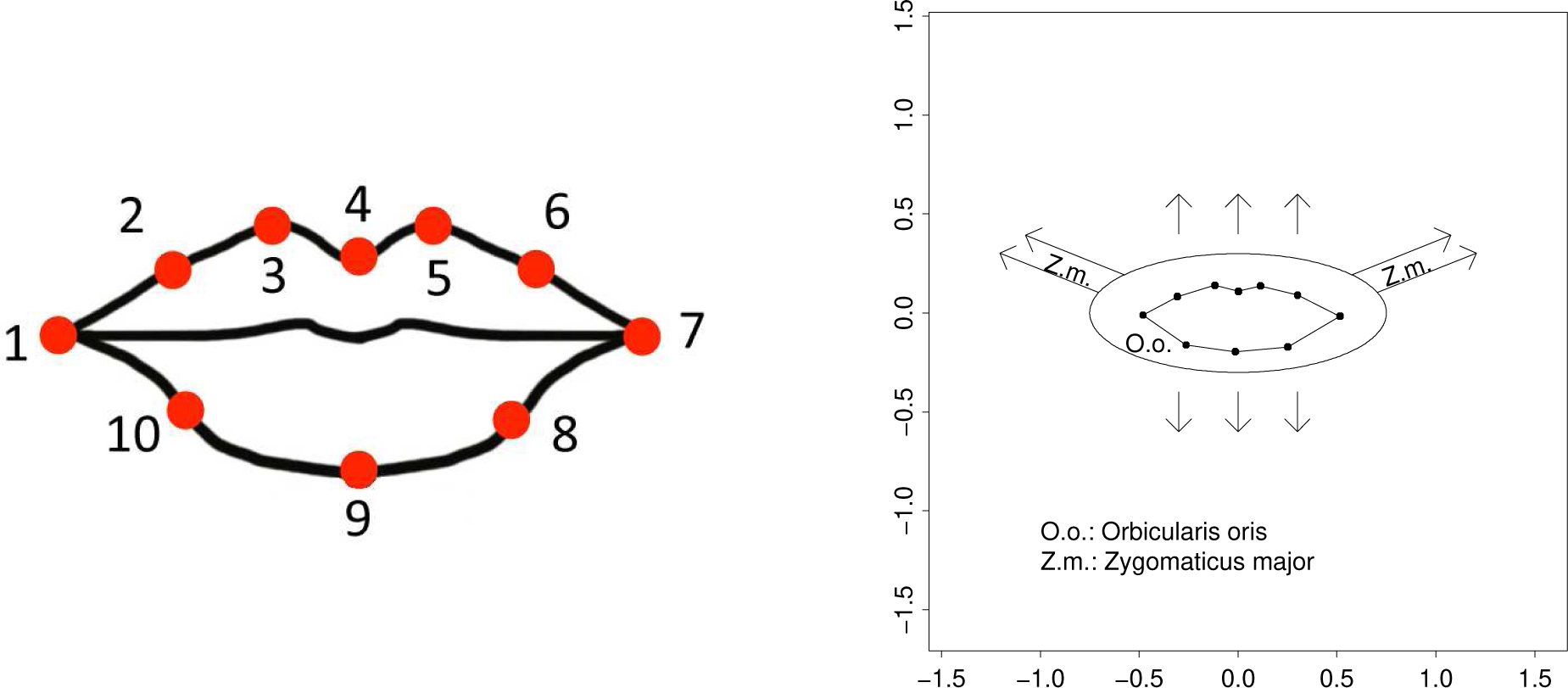
(left) A decagon of ten landmark points located on the vermilion border of our 14 subjects over two smile image sequences. Clearly a tremendous amount of information has been lost in the course of this informatic transformation. The purpose of the analysis in Section III is to hearken back from the decagon to the underlying anatomy anyway, by statistical manipulations. (right) Sketch of the two muscles, orbicularis oris and zygomaticus major, that appear to align with the geometric factors emerging from our morphometric analysis. For an anatomically appropriate version of this figure in its proper context, see any anatomy atlas, for instance, Gilroy and MacPherson, eds., 2017, page 516 or the cover (but see also Bookstein 2017b).

The contemporary literature of these muscle actions, as best one can tell from PubMed searches, focuses on their *reception,* i.e., their role in communicating emotional states. See, for instance, Niedenthal et al., 2010. This is also the central topic of a rather large literature retrievable under the rubric of “facial electromyography” that appears to treat these muscles as effectors evolved or at least intended to elicit such perceptions. Other specific foci of the facial muscle literature include the relation of orthodontic interventions to perceptions of the “attractiveness” of these muscle actions (e.g., the extent of visible incisors in the open-lip smile, the shape of the intercommissural line in the closed-lip smile) and the effect of a topologically significant birth defect, cleft lip, on all these aspects, along with the effect of diverse surgical repairs. The combination of these varied concerns renders the dynamics of the ordinary human smile, including the statistics of its comparisons, a worthwhile testbed for extending our methodological expertise in matters of the dynamics of repetitive motions of soft tissue. This article is intended as a contribution to that larger enterprise.

Some of this prior literature invokes methods of geometric morphometrics (GMM). These earlier publications include Ju et al., 2016, Darby et al. 2015, Hallac et al. 2017, Suhjaat et al. 2014, Campbell et al. 2012, and Johnston et al. 2003. But these and other more distant applications have not tapped the full power of the GMM toolkit. As sketched in the right column of Figure 2, these previous attempts have mainly relied on analyses of all the Procrustes shape coordinates as one extended list of variables analyzed landmark by landmark. This is not a good way to carry out analyses of forms that are as strongly integrated as these facial surface sequences are (Bookstein, 2015a) by virtue of their dependence on a short list of discrete muscle actions. In general it is considerably more powerful to analyze soft-tissue dynamics, and also to study group differences in dynamics, by converting the familiar Procrustes shape coordinates to a different set of basis vectors: the *partial warp scores* introduced by Bookstein, 1989, 1991, and briefly reviewed in Bookstein 2017a, 2018 and also here. (For an additional morphometric tool of a different sort, this one not yet part of the standard GMM toolkit, see Faraway and Trotman, 2011. We believe their approach is not pertinent to our data for the reasons given at our discussion of Figure 19 below.) While for a ten-landmark data set like this one the total count of partial warp scores is 2 × (10 − 2) = 16, we will ultimately argue that an analysis adequate for the present application involves only two of these scores.

### II. Data

#### The experiment

The data resource for the present study originated as a set of 3D motion capture records for 14 volunteer subjects (11 women, 3 men) producing and then relaxing each of two specified facial expressions named here by way of the Facial Action Coding system (Ekman et al., 2002). The closed-lip smile is action 12 of the FAC, hence “FAC12”; it is known in advance to be principally a contraction of the zygomaticus major. The open-lip or “maximal” smile combines this action with action 25 of the system, which is mainly a relaxation of orbicularis oris: hence, “FAC1225.” The teeth remained in occlusion over the course of all the experimental smiles. These two actions were selected out of the 32 in the current version of the FAC primarily because the corresponding smiles are among the facial actions considered among the most reproducible (Popat et al., 2010). All these smiles were *calibrated,* meaning that subjects had trained themselves (in mirrors) to extend the commissures while explicitly controlling the closure or not of the lips.

Of course this is only a stereotyped way of speaking. The corners of the lips are moved by other muscles than zygomaticus major, for instance, risorius and zygomaticus minor. Likewise, the opening of the lips is a consequence not only of the relaxation of orbicularis oris but also the contraction of levator labii superius and depressor labii inferius, two muscles that insert into the orbicularis oris. For the general role of a list of facial muscles in producing a variety of facial expressions, not only smiles, see Parke and Waters, 2008, or Daoudi et al., 2013. We are not aware of any earlier attempt to model the explicit dynamic variability of these facial expressions over individuals. For the related task of expression classification per se, see, for example, Pantic and Rothkrantz, 2000.

#### The ten (semi)landmarks

Our subjects’ smiles were captured dynamically as stereophotographs at a rate of 60 Hz over a time interval encompassing the full performance cycle and then were fused into a coherent surface mesh. Initial frames were oriented to the conventional anatomical axes (Frankfurt line horizontal, midsagittal plane aligned exactly forward), an orientation propagated through the rest of each series. Similarly, landmark points were placed manually on the initial mesh and then *they* were pushed forward automatically over all later surfaces of each sequence. For the purposes of this reanalysis the landmark location data were reduced to the two Cartesian coordinates in the (reconstituted) frontal plane only. Thus whenever we refer here to displacements of the lip commissures “upward and outward,” they are also being displaced *backward,* along a coordinate that we have chosen to ignore for the purposes of this particular data analysis.

Image sequences were truncated by one of us (BK) to exclude frames obviously prior to the onset of the expression elicited or obviously subsequent to the return to a resting state. There resulted a total of approximately 3000 individual frames. On the first frame for each subject, he marked the locations of the ten landmarks indicated in Figure 5. Four of these points are true three-dimensional landmarks in the sense of Bookstein, 2014: intersections of curves with other curves or with a local plane of symmetry. These are the outermost points 1 and 7, the *cheilions,* and points 4 and 9, *labiale superius* and *inferius,* defined as points of local bilateral symmetry along the upper and lower vermilion borders. The other six points are *semilandmarks* in the sense of Bookstein, 1991. Points 3 and 5 are the *christa philtri,* peaks on the lower margin of the philtrum, and the remaining points 2, 6, 8, and 10 lie midway along their curve segments above or below the midpoints of the corresponding chords. All these definitions apply only to the first image in any video sequence; landmarks in later frames were located by optical flow forward based on texture information local to the first video frame. For convenience we will refer to all ten of these points as “landmarks” in the text to follow, and to their net configurations as “decagons.” This particular decagon is a variant of that used by Darby et al. (2015). We have replaced their landmarks stomion and sublabiale, which do not lie on the outer vermilion borders, by the new upper lip semilandmarks 2 and 6, thereby covering the template in Figure 5 more evenly.

The textured surface data underlying these landmark locations were collected using a 3D motion capture system (4D) marketed by Dimensional Imaging Ltd, Glasgow, UK. The tracking of pixels from one image frame to the next was by DI4DView software, which offers the subpixel precision necessary to follow landmark points forward in time. The ten-landmark configuration of homologous points around the vermilion border was followed over a two-second video at 60 Hz for each of two smiles for the 14 subjects in this study. The original data are in three dimensions, not two, and the original list of landmarks includes four others much higher on the head that were used to supply a fixed coordinate system for earlier analyses. The black lines in the left-hand panel of Figure 5 are only to aid in visualization; they are not a component of the measured data.

#### Data reduction

Because the expressions under study extended over varying intervals of elapsed time, each was reduced to a sequence of precisely 100 intervals (101 configurations) by linear interpolation within the limits of the truncation. Figure 6 shows the sample averages of these motions as originally captured. (Each composite sequence was centered around the average of its 10 landmarks over the full set of 101 stages of the motion, the coordinate system already introduced in the central column of Figure 2.) The count of original frames varied from 91 to 157 over the full sample of 28 smiles. Figure 7 suggests the quality of the optical-flow tracking for a sample of three frames for the open-lip smile of subject 12. Figure 8 shows the complete record of 101 resampled lip shapes for this single open-lip smile as projected onto the best conventional representation plane, the first two relative warps (principal components of shape) of that full sequence. The smoothness here testifies to the basic stability of the video system, as does the very small amplitude of the “vibration” seen at the ends of the trajectory, the time intervals during which the subject was to hold either the achieved smile or the relaxed face fixed over several frames. Notice, however, that the sequence is not a single trajectory traversed twice — the relaxation of the achieved smile is not the reverse of its implementation. Indeed this cycle is not even a closed loop: the final rest position is slightly displaced from the original rest position.

**Figure 6.**
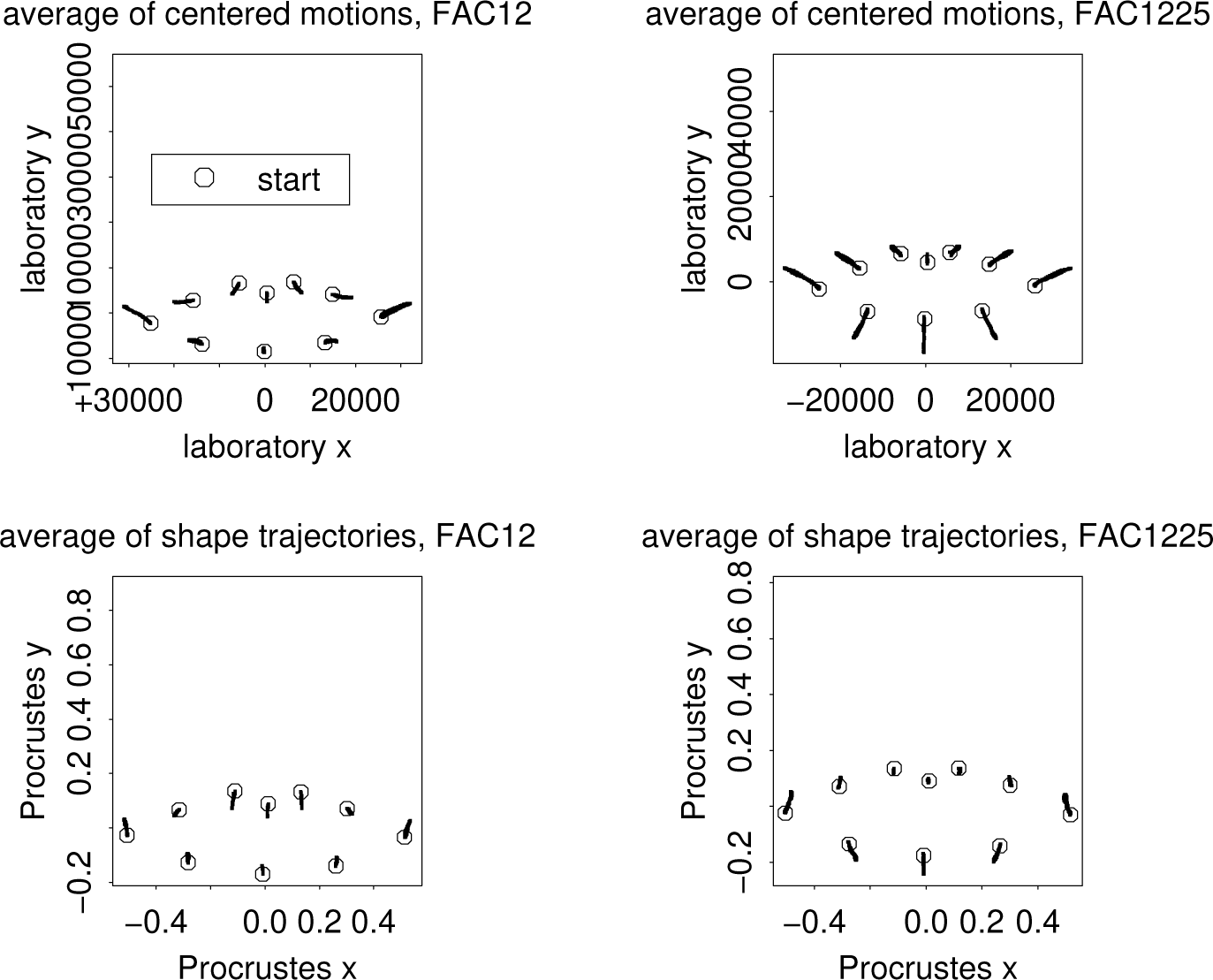
Results of the motion capture in frontal view: Average trajectories of these 14 pairs of smiles. (top row) In centered laboratory coordinates, units of microns. (bottom row) The same after Procrustes registration, which for these data involved only size standardization, with hardly any rotation. Open disks: averaged starting positions.

**Figure 7.**
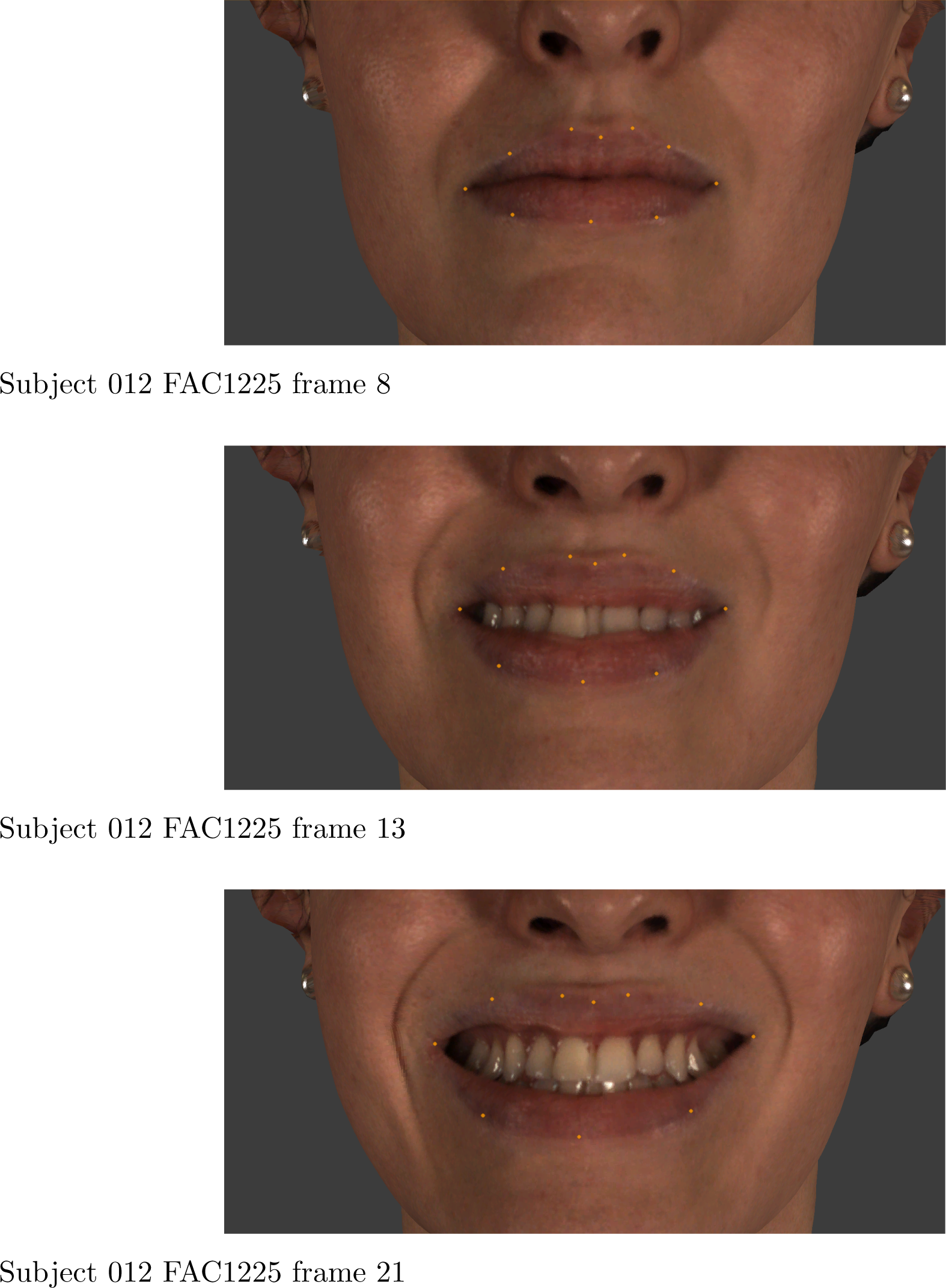
Three frames from the original frontal video for subject 12, whose FAC1225 smile is the example in Figure 8. The three frames here correspond to the three spacetime landmarks (beginning frame, solid dot, ending frame) on the curve in the second column of the third row in Figure 19.

**Figure 8.**
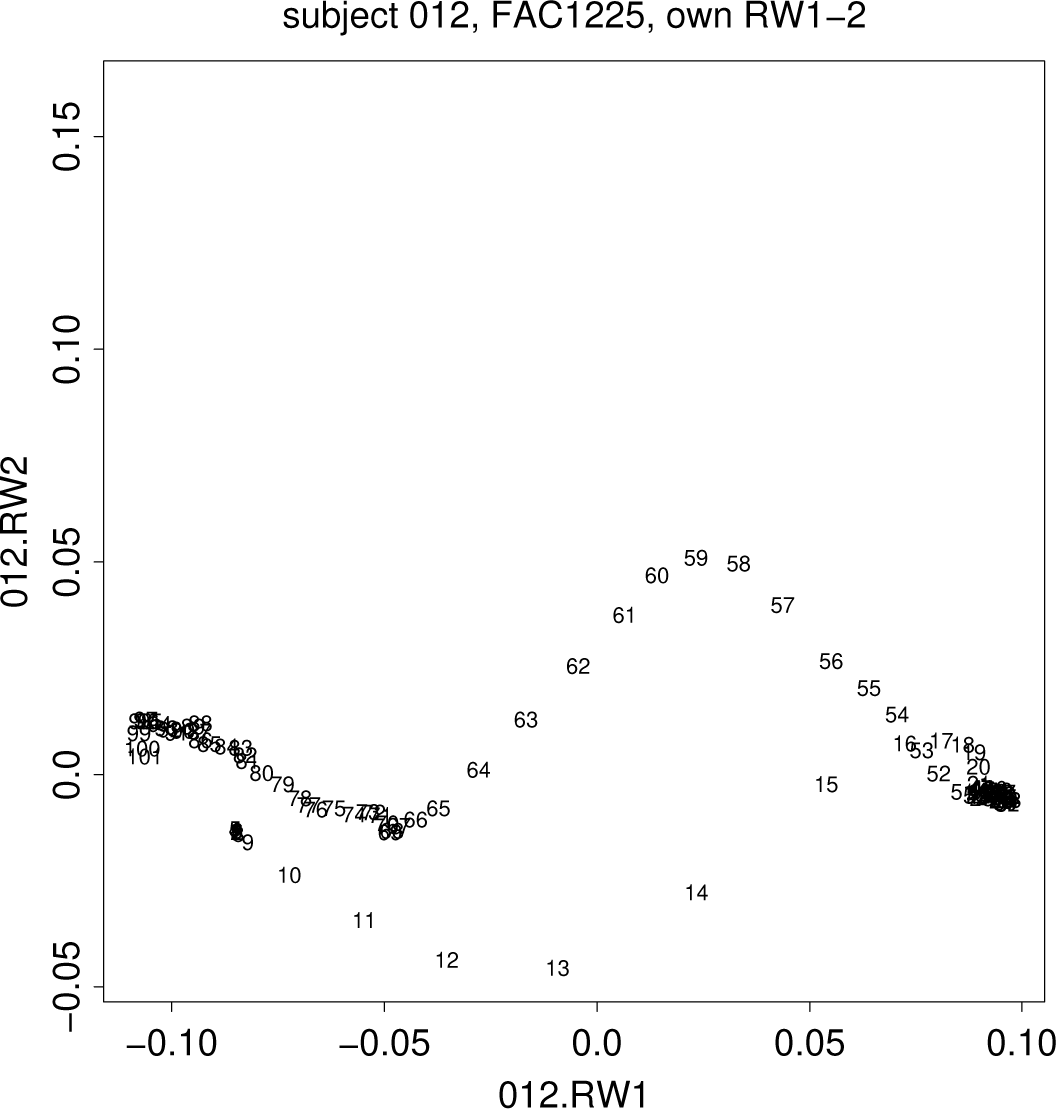
Reliability of these representations. Shown here is the plot of the first two relative warps (principal components of the Procrustes shape coordinates) for this same subject over the full 1.57-second data record. Numbers are resampled frame numbers from 1 to 101. Later figures for this subject in this condition involve the first 33 of these frames. See text.

### III. Morphometric methods and empirical pattern analyses

#### Decagons of shape coordinates

Our analysis begins with the tableaux in Figures 9 and 10, which present the complete shape sequences for each of the 28 smiles examined (two smiles for each of the 14 subjects), each one after its own separate transformation into the standard linearized Procrustes shape space at right in Figure 2. These are the only data resources used for any of the computations to follow — all other diagrams are simplifications of the information in this pair.

**Figure 9.**
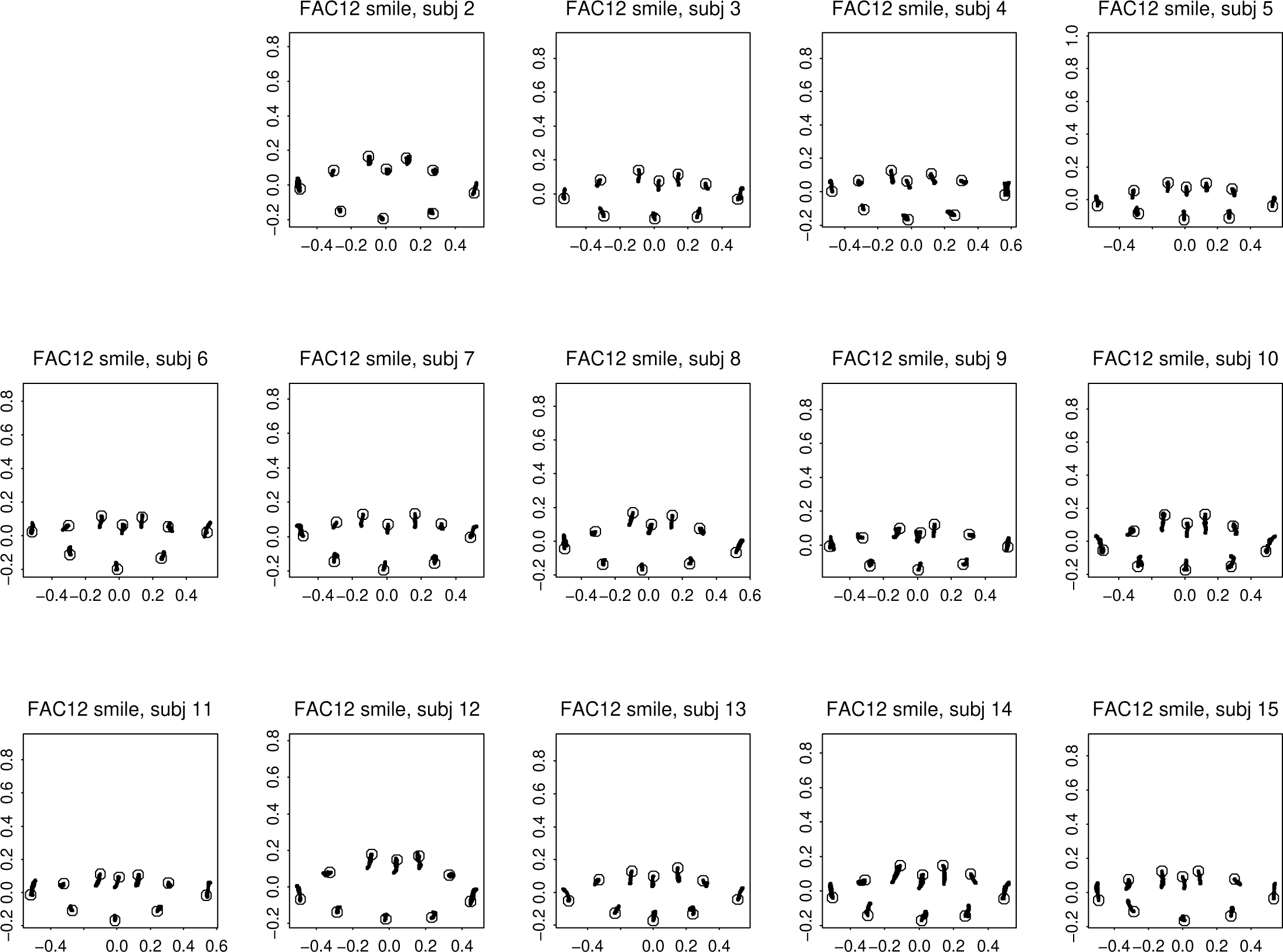
Complete Procrustes records for the FAC12 (closed-lip) smile for each subject separately, resampled at 101 evenly spaced interpolated times between subjectively set moments of stasis prior to initialization of the smile’s dynamics and following the cessation of additional movement. The large circles indicate the initial rest configuration. Subjects are numbered from 2 to 15 in order to preserve their original numbering, which included a subject 01 whose data were incomplete.

**Figure 10.**
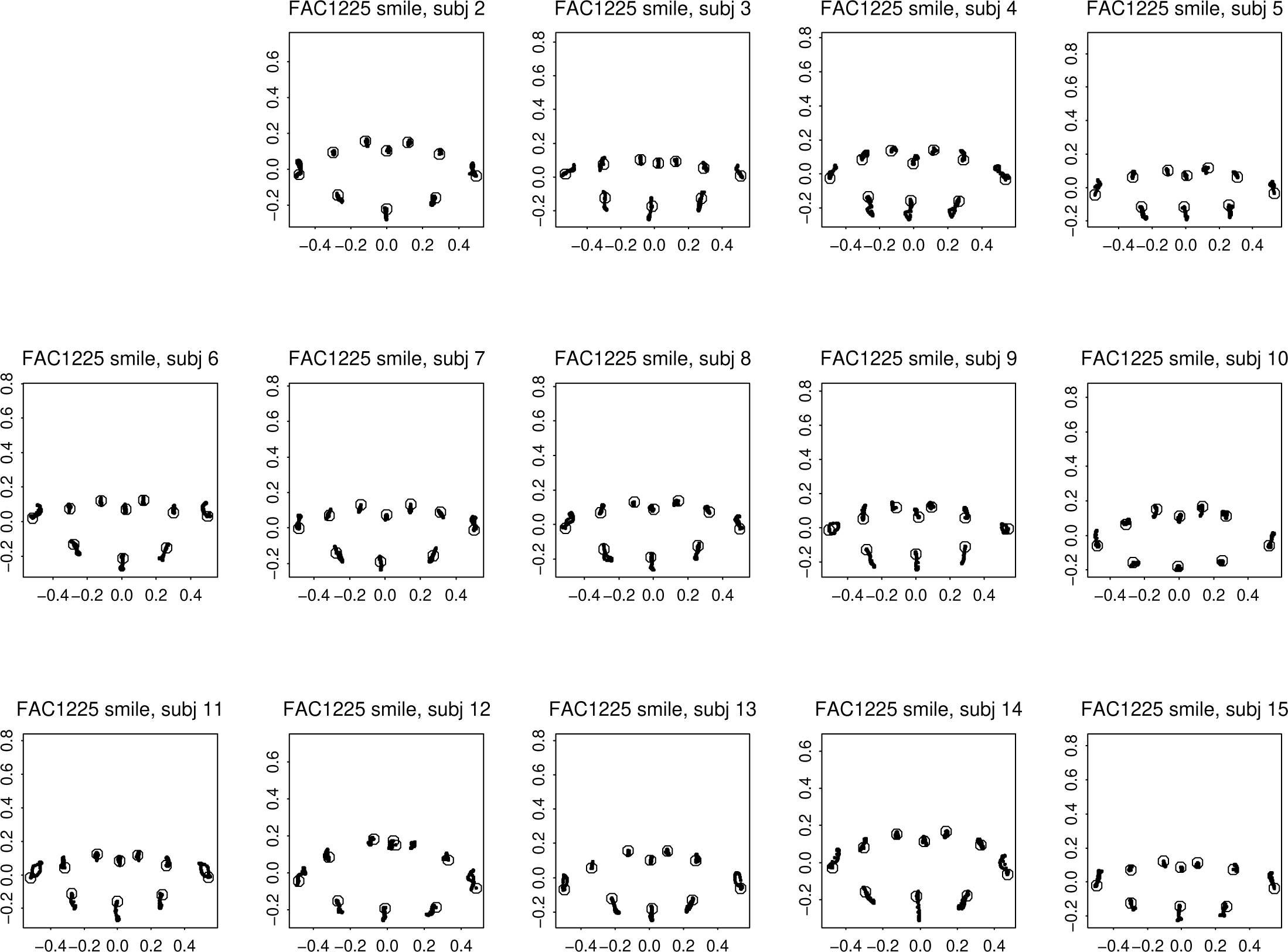
Analogue to Figure 9 for the FAC1225 (open-lip) smile.

The obvious difference between the two scenarios in the bottom row of Figure 6 persists through panel-by-panel comparisons of Figure 9 to Figure 10. In both smiles the outer corners of the lips typically appear to shift upward relative to the center of “mass” of this outline: a little bit outward for the closed-lip, a little inward for the open-lip in view of the increase in decagon height in this condition. But in the closed-lip smile (Figure 9) it is the landmarks of the *upper* lip that shift relatively downward in compensation, whereas in the open-lip smile (Figure 10), with the exception of subject 10, it is the landmarks of the *lower* lip that shift their relative positions in this way. Note (especially in panels like that for subject 11 in Figure 10) that, as in Figure 8, many of these are loops — the sequence of frames along the relaxation does not duplicate those during the creation of the smile.

#### Smile-by-smile relative warps

Plots like these, however drastically simplified they are in comparison to the original anatomical setting, nevertheless still preserve too much information to serve *per se* as part of a biometric analysis. We begin our pursuit of an appropriate pattern description by reducing each shape trajectory to a sequence of points along a projection onto a specific plane within their 16-dimensional shape space: the plane specified by the first pair of relative warps. Recall that this is the plane within this linearized space having the greatest central moment (sum of variances along any two perpendicular axes). This can also be thought of as the plane of the first two principal components of the 14 × 101 = 1414 shape coordinates pooled over all the frames in Figure 9 (for the closed-lip smile) or Figure 10 (the open-lip smile). The resulting displays are set out in Figure 11 for the decagons derived from the 14 closed-lip smile videos and in Figure 12 for those from the 14 open-lip smile videos. We further unclutter the figures by truncating each sequence at the moment of maximum displacement along the horizontal axis, the best pooled estimate of “the net change caused by the smile” in this shape space. Now that there is room to number individual frames, we can also trace the *pace* of these smiles, the ramping-up and then ramping-down of the speed of these deformations as the responsible muscles begin their action, move to a brief steady state, and then slow to a new stasis (subjects were instructed in advance to “hold the smile” for a little while).

**Figure 11.**
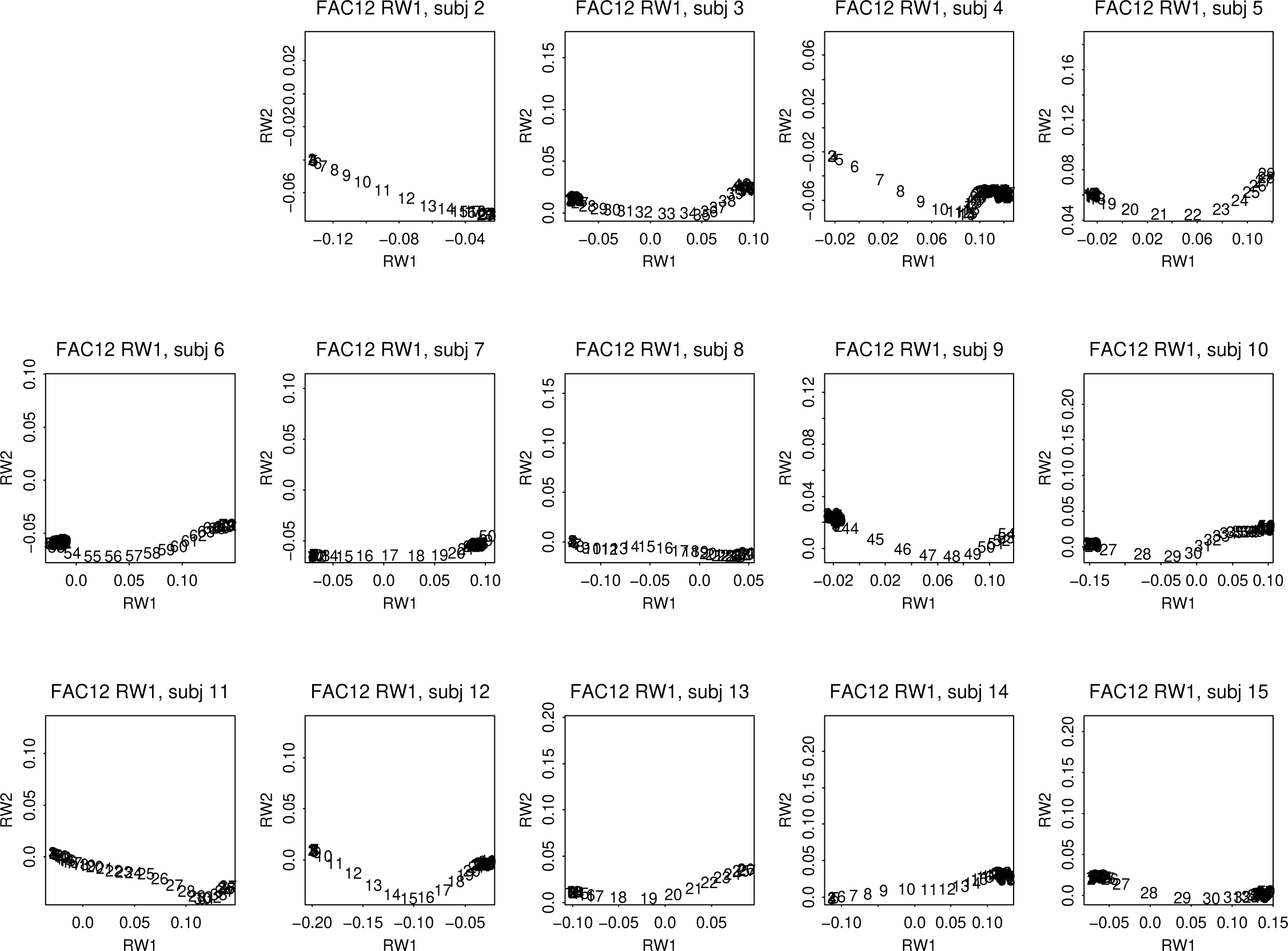
Subject-by-subject plots of the projections of the series of landmark configuration shapes on the first two relative warps (Procrustes principal components) of the pooled data set of 14 × 101 = 1414 such configurations for the closed-lip smile. Each sequence of plotted points has been truncated at the moment of maximal extension along pooled relative warp 1. In these plots, integers are resampled frame numbers (between 2 and 101) corresponding to the normalized temporal position of each configuration within the full cycle of the smile that was elicited. Counts of frames remaining after the truncation range from 24 to 72.

**Figure 12.**
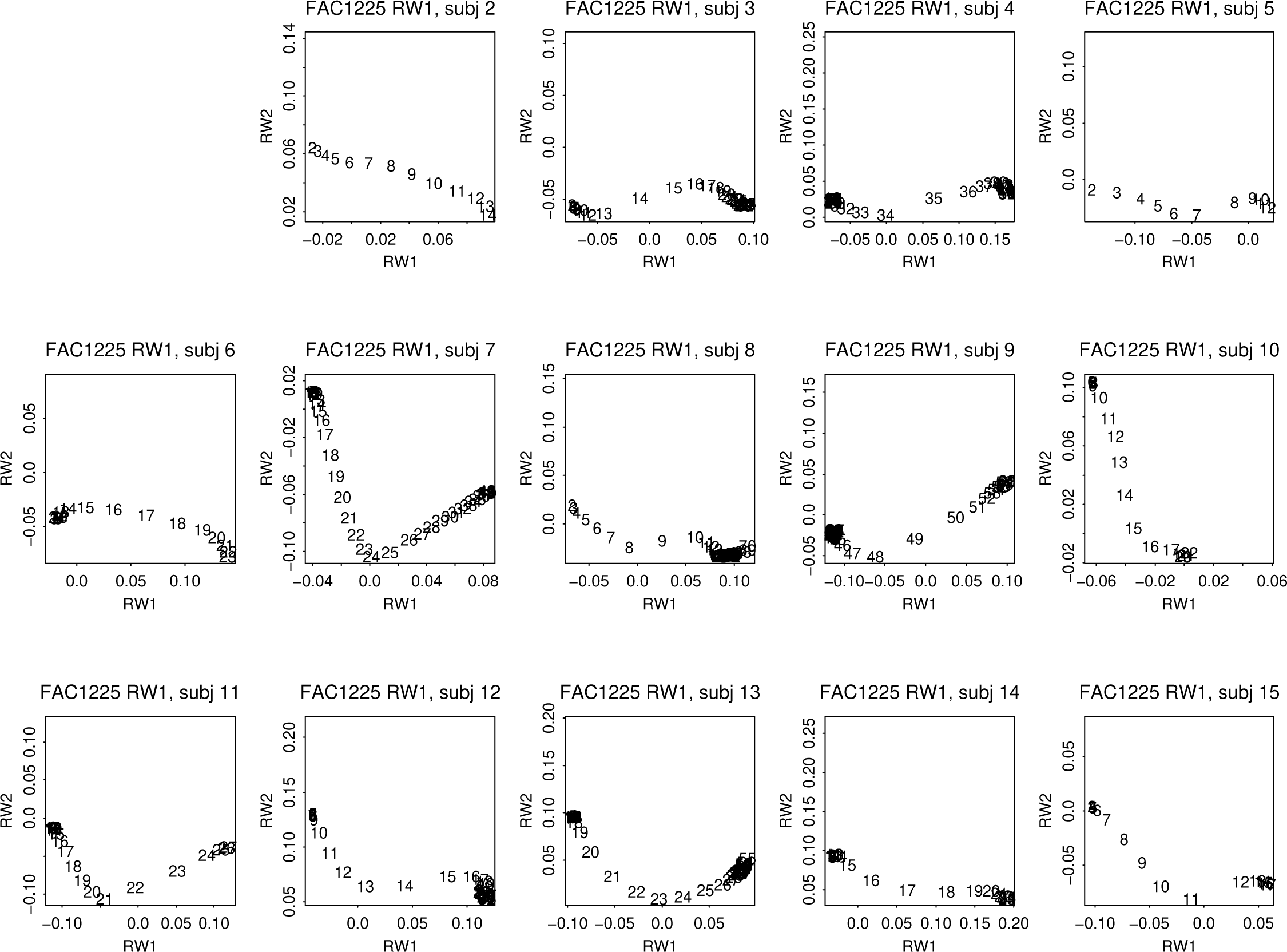
The same subjects’ open-lip smiles displayed using the same convention. Counts of frames remaining range from 12 to 74.

The dynamics of the series in Figure 11 can reasonably be construed as a homogeneous bundle of monotone progressions along nearly straight paths the curvature and angulation of which vary modestly from subject to subject. In the modeling phase of our work it will be represented as the expression mainly of a single factor whose orientation varies similarly modestly across subjects. We will argue that this corresponds well to the physiologist’s conception of the action of a single muscle or singly innervated group of muscles across such a sample of normal humans. The corresponding count of parameters is two, corresponding simply to the vector from end to end of the trajectory. We do not claim that progress along these trajectories is linear, sigmoid, or of any other specific functional form. (Variations of curvature will be considered briefly in the next section.)

But the same claim of underlying homogeneity cannot be made about the considerably more diverse trajectories in Figure 12. Considerable variation is evident in the visual impact of these curves. While a few of them are still approximately flat from left to right, others show substantial evidence of a two-phase structure, progress that is initially linear in one direction (downward on these plots) but is then redirected in an upward or horizontal direction. One might say that this bundle of trajectories is heterogeneous, or, better, that it represents a superposition of two factors per subject of which the second may be triggered later than the first and (in most cases) runs in a direction of shape space oblique to the first.

### On integration: only the two dimensions of largest scale need to be examined

From this collection of familiar GMM graphical maneuvers we turn to one that is likely not so familiar: the analysis of *integration* introduced in Bookstein, 2015a, and revisited in Bookstein (2015b, 2016, 2017a). This style of plot was developed to summarize the geometric scaling of shape-coordinate distributions for a variety of comparative purposes. As before we offer you two multipanel displays, one (Figure 13) for the fourteen closed-lip smiles and the other (Figure 14) for the fourteen open-lip smiles of the same subjects.

**Figure 13.**
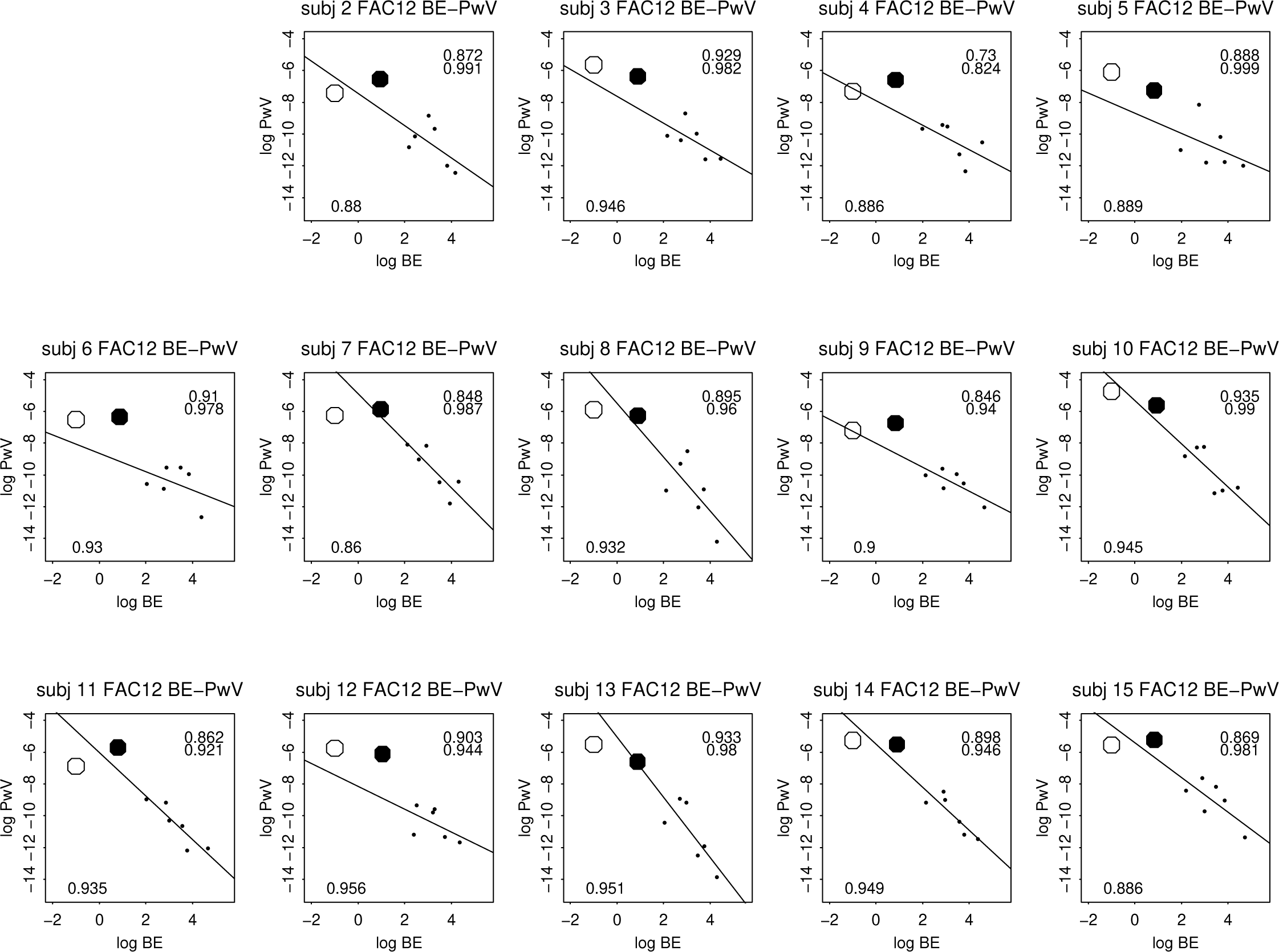
Bending energy–partial warp variance (BE–PwV) plots for the closed-lip smiles of the 14 subjects. Line segments are for regressions on the six small dots, omitting both the large filled dot (for PW1, which does not come under the residual analysis here) and the large open circle (the uniform component, which does not actually have a horizontal position). Tick marks on the axes here and in the next figure are spaced at multiples of *e*^2^ = 7.39. See text.

**Figure 14.**
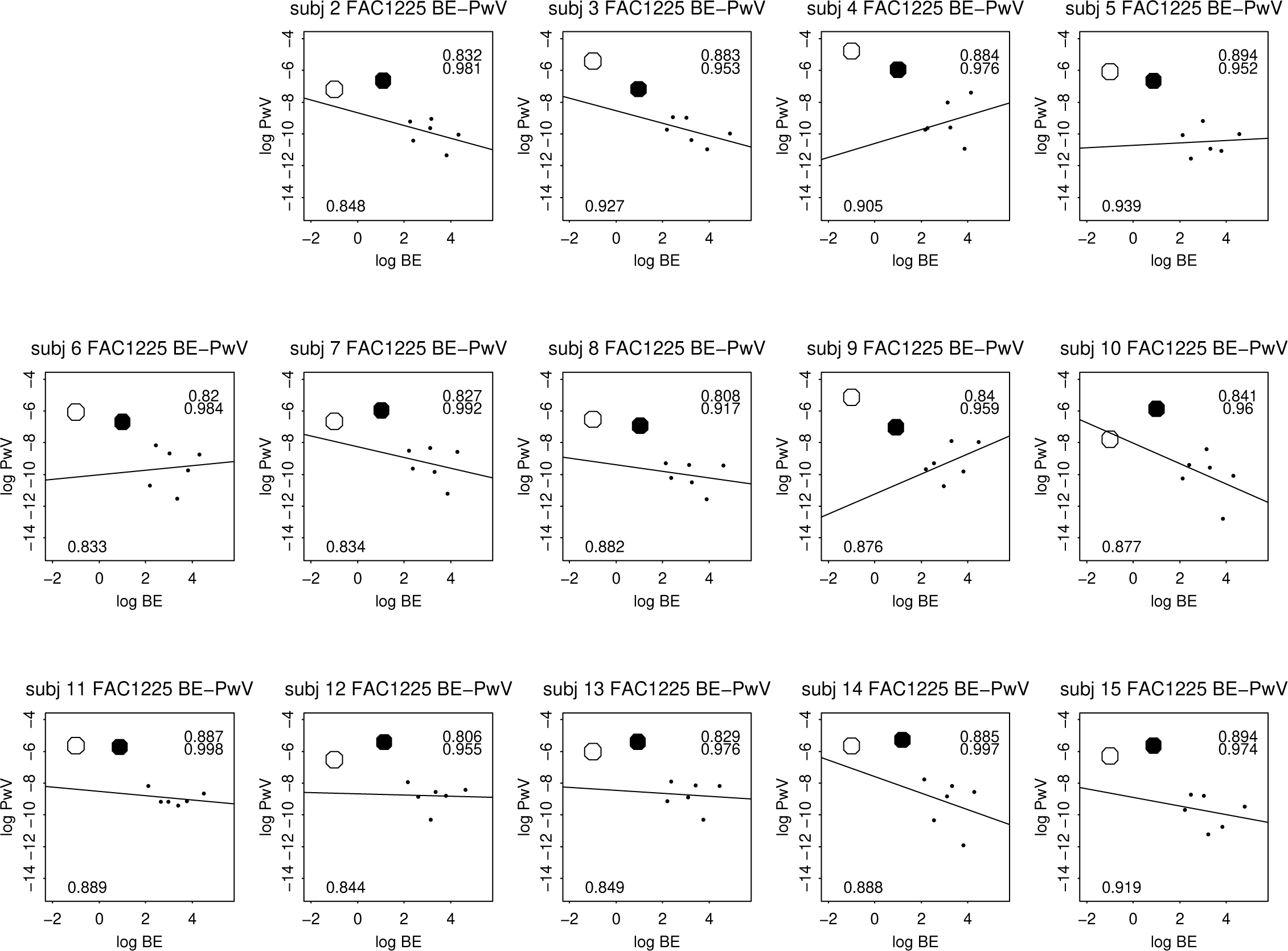
The same for the open-lip smiles for the same fourteen subjects.

In both Figure 13 and Figure 14, the vertical axis is for log-transformed summed variances of the horizontal and vertical components of the partial warps that decompose any landmark shape distribution around an average into a sum of eigenfunctions of the bending energy matrix of that mean shape. The horizontal axis in these plots combines a value of −1, arbitrarily assigned to the uniform (affine) term in this decomposition — its actual bending energy, zero, does not have a logarithm — with the logarithms of the seven actual nonzero eigenvalues of these eigenfunctions, thus a total of 8 log-variances (the landmark count, minus 2) for these decagons. The leftmost two points are plotted with large symbols because they will prove to bear the most signal according to the plots of relative warp loadings in Figures 15 ff. That for the uniform component is hollow because its *x*-coordinate is arbitrary. Those for the partial warps after the first are plotted with smaller dots because they invariably explain less shape variance.

**Figure 15.**
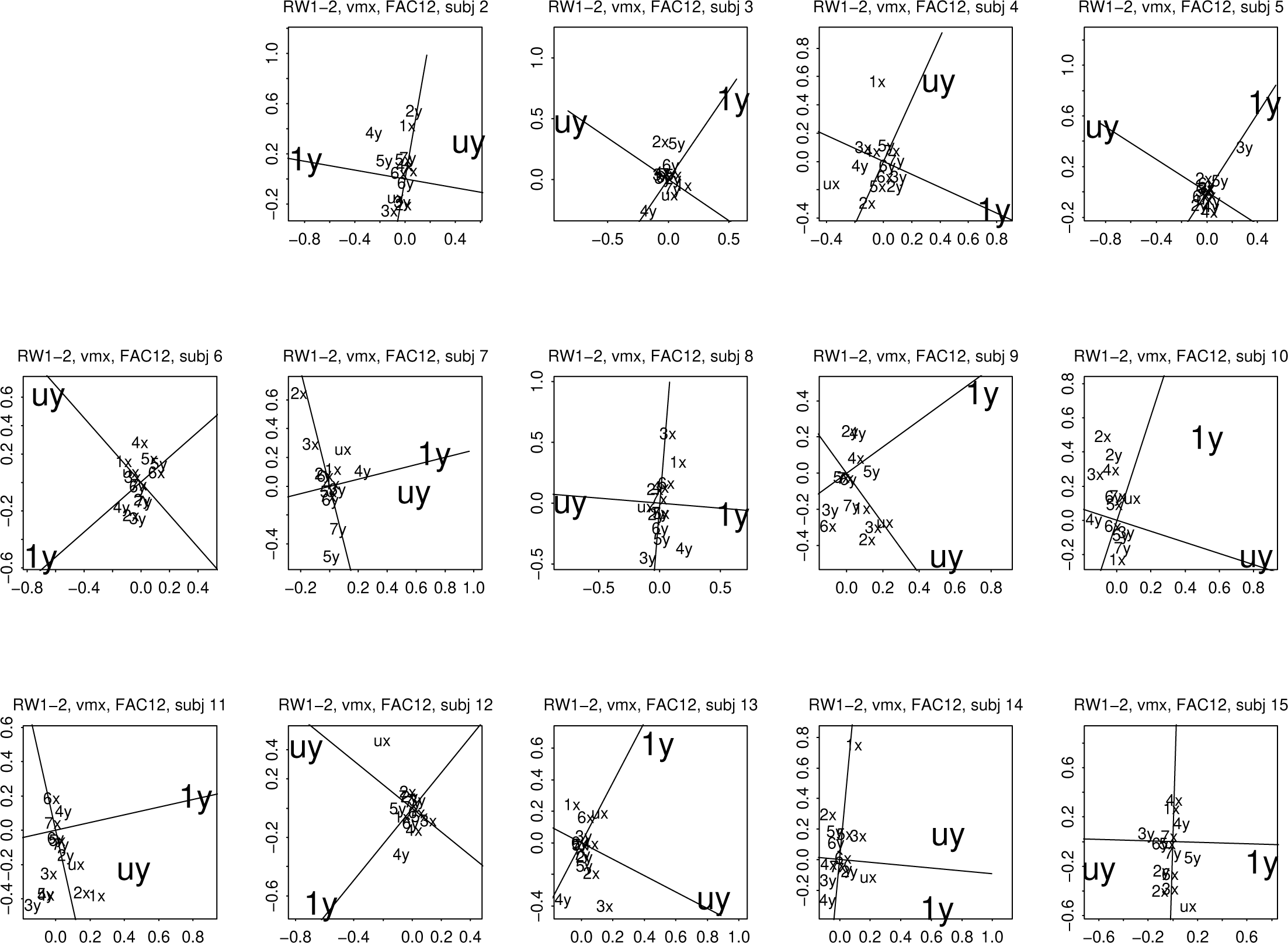
Scatterplot of loadings on the first two relative warps of the opening trajectories for the closed-lip smiles, Figure 11, subject by subject. These are the loadings of a relative warp analysis to the basis of the 16 partial warp scores. Light lines at 90◦ to one another: varimax rotation of the original abscissa and ordinate. See text.

The straight line in each panel is the ordinary linear regression through these six small dots. The slopes of these lines vary from −1.9 to −0.64 around a mode of about −1. These are all in the range that Bookstein (2015a) called “self-similar,” the regime previously analyzed from a more mathematical point of view in Mardia et al., 2006. In this range, variance of a given shape (of a real or interpolated landmark subconfiguration) is invariant to the geometric scale of that subconfiguration, corresponding to an approach to a simple instrinsic random process model on the lip outline for the residuals of the original landmark locations after partialling out those two patterns of largest geometrical scale. A formal model for these 14 shape sequences would do well to begin with a combination of two subject-specific parameters, one for each of the factors underlying the pair of large plotting symbols.

The decimal quantities printed on these panels are as follows. In the lower left corner appears the fraction of total subject-by-subject shape variance that is accounted for by the first two partial warps, PW0 and PW1; in the upper right corner, the fraction of the total variance that is accounted for by just the two dimensions of symmetric change, vertical components of those same two partial warps; just below, the ratio of the upper-right sum of squares to that at lower left. All these ratios are typically near or beyond 0.9 throughout this sample, reassuring us that the simplified pattern language at which we eventually arrive exhausts nearly all of the geometric information in these data.

Figure 14 presents the same 14 panels for the condition of the open-lip smile. Now the regressions of log variance on log bending energy for the six smallest-scale eigenfunctions range in slope from −0.64 to +0.64, thus never varying far from zero, which is the value they would assume were the residuals from the combination of the uniform term and this largest-scale partial warp actually distributed as independently digitized points of the same digitizing variance at every point in every direction (Bookstein, 2015a). The reduction of these trajectories to the two-dimensional model we will introduce presently is thus even more strongly supported by the empirical pattern of shape variances over geometric scale in this second of the two experimental conditions.

### Varimax factor analysis confirms the bidimensionality

The predominance of the same two partial warps, PW0 and PW1, in all 28 of these smiles suggests a potential route toward a powerful and unexpected statistical summary. Figures 15 and 16 together present the usual set of 28 replicate panels for an analysis that expresses the relative warps of each smile *separately* in terms of its own suite of 16 partial warp scores. In each panel, all the partial warps are identified separately as *u* (uniform) or by the integers 1 through 7; the two Cartesian coordinates of each supply the suffixes *x* and *y*. The loadings for *uy* and 1*y* are printed in larger font because in nearly every frame they grossly dominate the underlying relative warp loadings. This can be confirmed by the *varimax rotation* of these scatterplots, still subject by subject, to the rotated coordinate system (new orthogonal axes) that maximizes the sum of the variances of the squares of the corresponding rotated loadings. These new axes (light orthogonal lines in the figure panels) often (especially for the open-lips smile, Figure 14) pass nearly through the loadings printed in large font, which, not coincidentally, usually lie at an angle of nearly 90^◦^ out of the center. That a varimax rotation to two factors works in this context is likely because of the dominance of just two muscles in the signal here. The general case offers no such convenience. For an explanation of the history of the varimax method and its pertinence to the analysis of partial warp scores, see the exegesis and the examples in Bookstein, 2017a.

> *Algorithmic note.* The method of varimax rotation has been cited over 6000 tines since its first publication by Kaiser (1958). It was previously recommended for applications in morphometrics by Reyment and Jöreskog in their textbook of 1993, but, to our knowledge, was not suggested for use in GMM applications prior to the announcement in Bookstein, 2017a. The Cartesian rotations diagrammed in these two figures were not computed by any public “varimax” software package. Instead, they were generated by the newly discovered formula explicated in Bookstein’s appendix.

**Figure 16.**
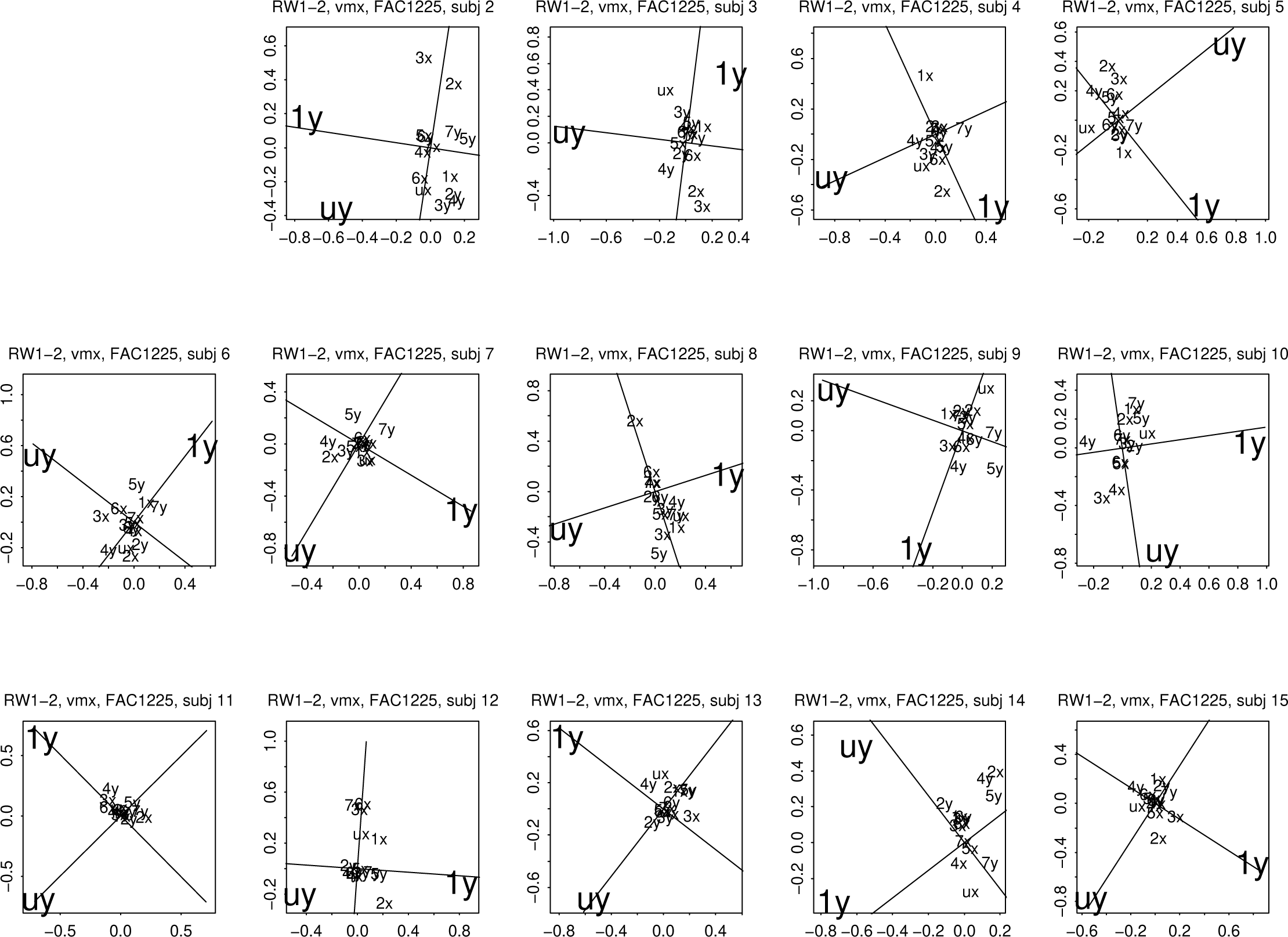
The same for the subject-by-subject open-lip smiles, Figure 12. The terms *uy* and 1*y* now dominate even more.

The net import of Figures 15 and 16 is striking: in all 28 smiles, the loadings of partial warps *uy* and 1*y* are by far the largest (confirming, in turn, the decision to base multivariate analysis on the partial warp scores instead of the shape coordinates individually). This impression is confirmed by the summary superpositions in Figure 17, which combine all subjects for each of the two smile conditions. The systematic dominance of just two partial warp features, the vertical component *uy* of the uniform term and the vertical component 1*y* of the largest-scale bending term, is now obvious. Quadrants of this plot are arbitrary inasmuch as the polarity of the axes is not comparable from subject to subject; the figure’s import is conveyed purely by distance from the point (0, 0) at the center. The convex hull of the (*uy,* 1*y*) pairs is a unit circle, not a rectangle, because the points of this plot are projections of loci originally residing on the unit sphere in their 16-dimensional space.

**Figure 17.**
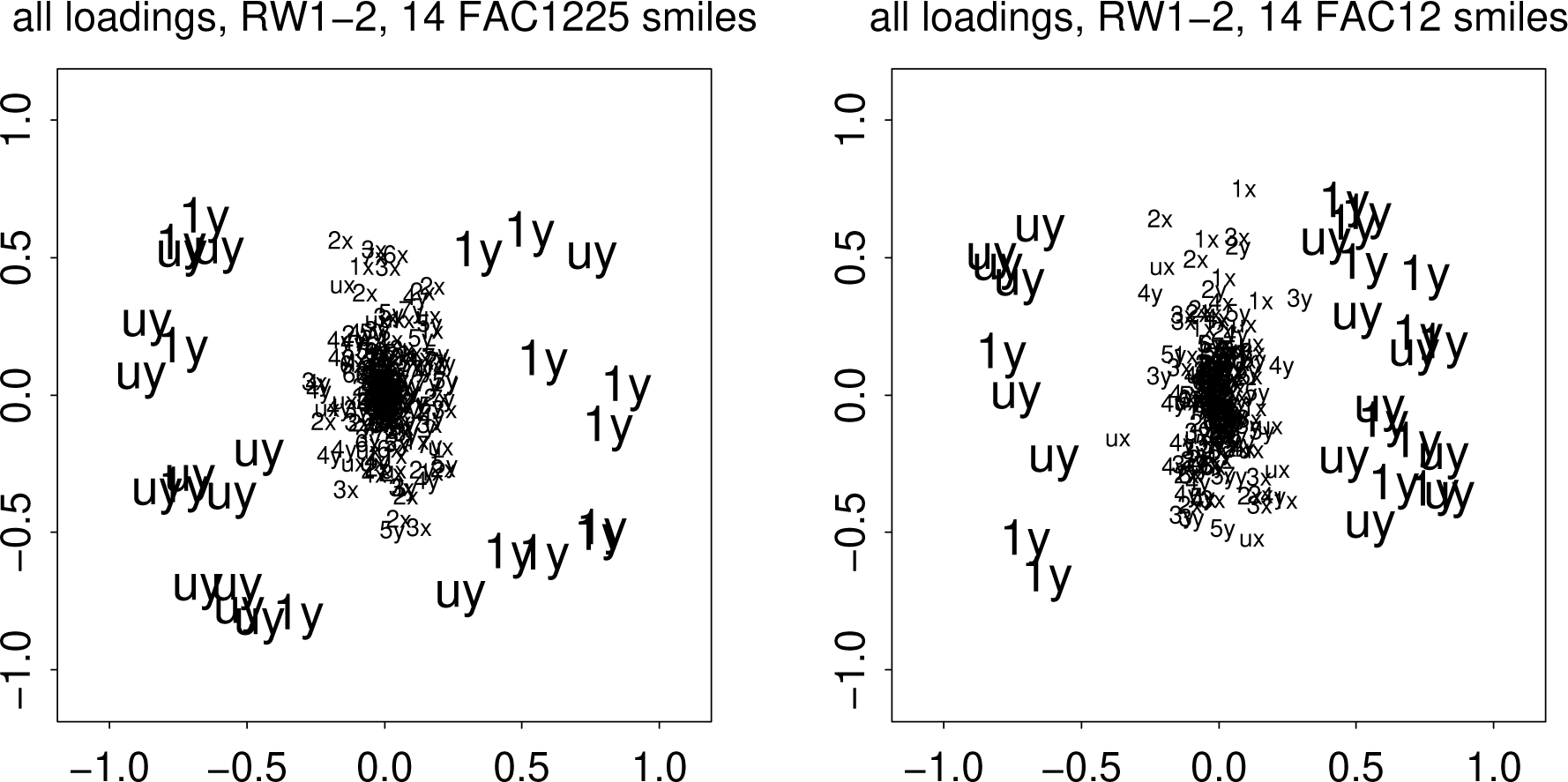
Superposition of all the frames in Figures 15 and 16, respectively. Each plot thus displays 14 × 16 = 224 partial warp loadings. Around the outside of either plot are a total of 28 symbols, 14 for *uy* and 14 for 1*y*. Inside are the other 196 loadings; they overlap too much to be readable.

#### The two factors can be identified with single partial warps …

We can highlight this finding further, therefore, by discarding all those small partial warps that piled up in the middle of the scatters in Figure 17, thereby reducing each smile trajectory to its projection just on these two components of largest amplitude. (These two components are now computed separately for each truncated smile for each subject.) The 14 × 2 = 28 separate projected smile trajectories are set out in Figures 18 and 19 for the closed-lip and open-lip smiles, respectively. This projection onto two dimensions is different both from the standard GMM projection onto the first two relative warps, Figures 11 and 12, and from the projections of Faraway and Trotman (2011) onto the straight line linking initial and extremal configurations. (The trajectories in this data set are clearly not straight.) The geometry of the renderings here will be diagrammed more explicitly in Figure 20.

**Figure 18.**
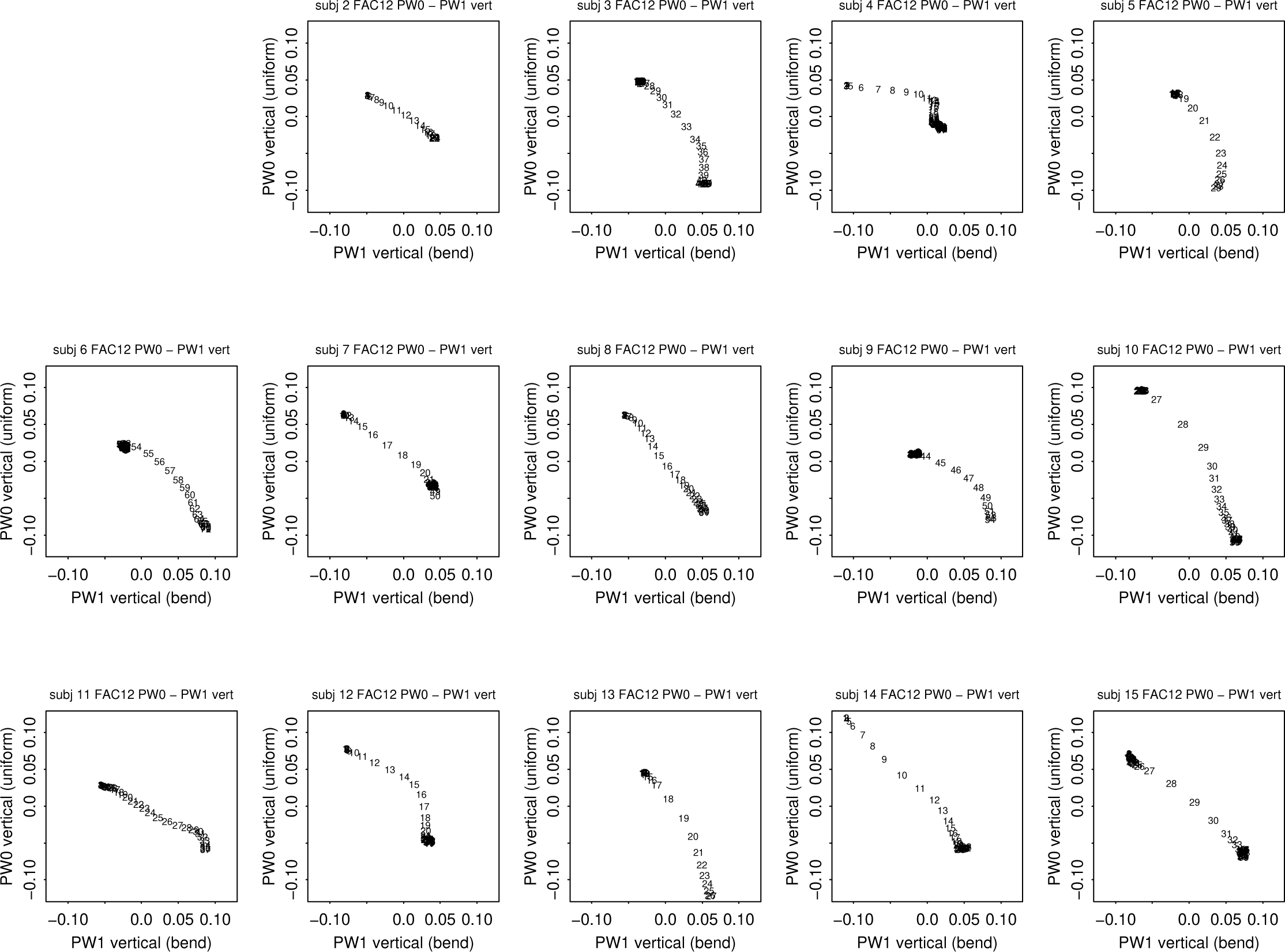
Projection of the closed-lip trajectories, Figure 9, onto the plane of two components only, *uy* and 1*y* as motivated by Figure 17. Vertical axis: vertical dilation (here, lateral extension). Horizontal axis: bending of this horizontal.

The majority of trajectories in Figure 18 (the closed-lip smile) are not too curved, and what curvature they have is always in the same direction (northeasterly here). This means that for those paths that are not straight, peak displacement rate along the bending component (horizontal) usually precedes that along the vertical component (closing) as encoded in the interpolated frame numbers printed. In Figure 19, however, most of the trajectories sustain a marked change of direction en route. In each panel a candidate for this change point has been marked: the point farthest from the end-to-end chord of its trajectory. At most of these extremal points, the direction of the trajectory changes sharply. Our physiological interpretation is that the onset of orbicularis relaxation lags the onset of zygomaticus contraction by that time interval in these cases.

**Figure 19.**
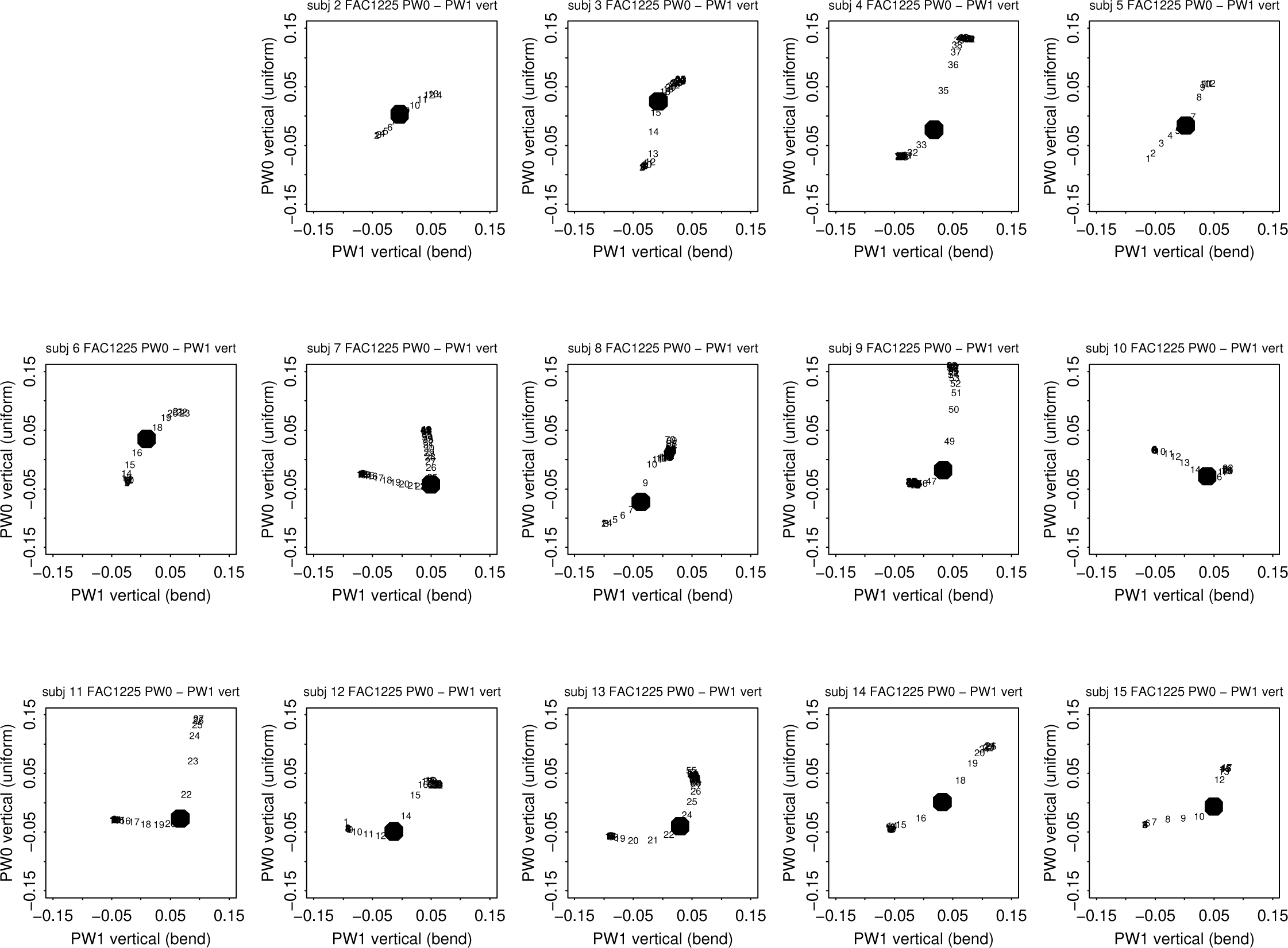
The same for the open-lip smile trajectories in **Figure 10**. The marked point in each panel is the one farthest from the end-to-end chord of its trajectory.

These figures, in turn, permit superpositions that are highly suggestive of the modeling approach we will shortly be pursuing. First, we superimpose all the trajectories of each condition so that they spring from the same starting point (the big solid dots in either panel of Figure 21). It is evident that the two smile requests lead to wholly different physiological responses, as conveyed by the endpoints (large open circles) of these same 14 paired smiles. The separation of the two types of smile is thus perfect, a highly desirable outcome in any experimental morphometric study.

**Figure 20.**
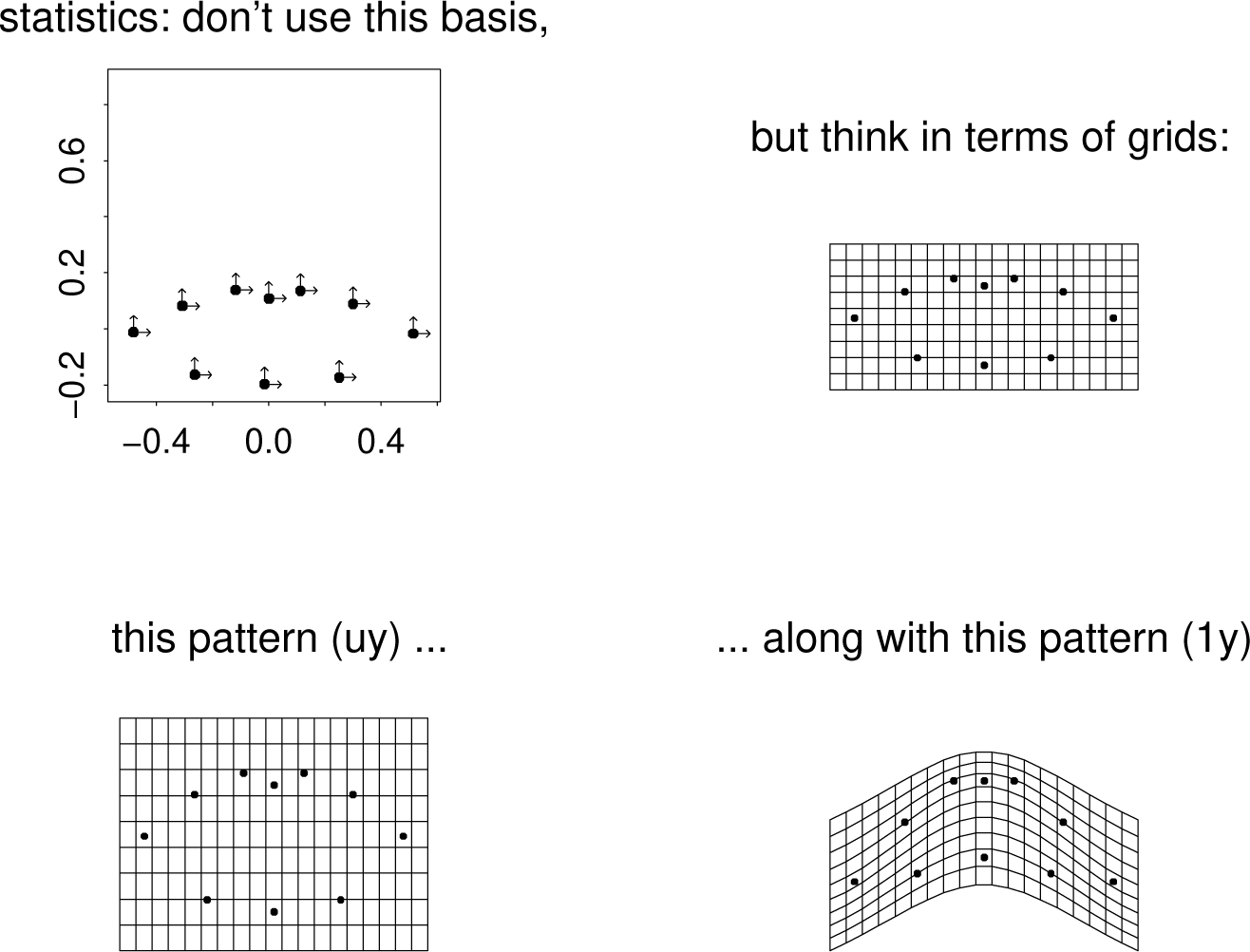
To achieve power in our GMM analysis of this integrated organismal phenomenon it was necessary, *as it usually* is, to switch from the näıve basis of the full set of Procrustes shape coordinates (upper left panel — there are 20 of these, corresponding to x− and y-coordinates of each of our ten landmarks) to an appropriate selection from the roster of partial warp scores just reviewed. These are most easily drawn as deformations of a grid (upper right) rather than pointwise displacements. Of the 16 partial warp scores that are required in order to exhaust the information in this ten-landmark data set, our smile analysis will ultimately involve only two, the vertical components of the uniform term (lower left) and the largest-scale bending (lower right), corresponding, we will argue, to the actions of the orbicularis oris and zygomaticus major muscles, respectively.

**Figure 21.**
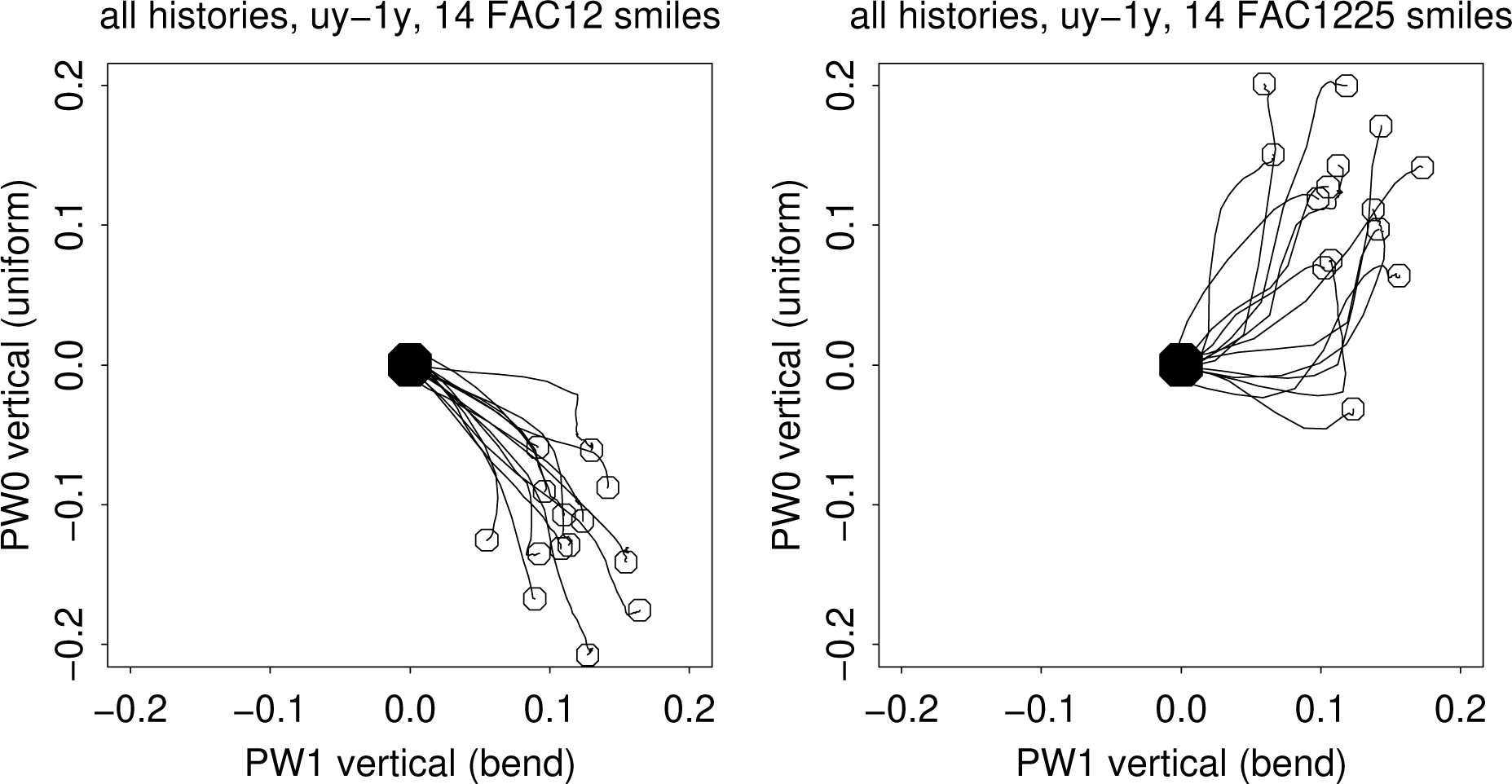
Summary of the projected trajectories of Figures 18 and 19 as superimposed over all 14 subjects as if all smiles began at the starting configuration plotted at (0, 0) here (the big black dot). Left, the 14 closed-lip smiles; right, the 14 open-lip smiles. Open circles, “final” state of the same smiles (state of greatest projected distance from the starting form along pooled relative warp 1), as previously displayed in Figures 18 and 19. At right, the lowest of these open circles is for subject 10, whose FAC1225 data are described in detail in the text. Horizontal and vertical axes, vertical bend and vertical dilation, again as in the preceding two figures.

It further appears that there are differences of muscular timing between these smiles. Regarding the horizontal coordinate (the bending) they begin similarly with an increase that is monotone in most cases, but while the closed-lip smile for most subjects involves a relative vertical compression almost immediately, that for the open-lip smile often delays the onset of that component of the shape change, which, with only one exception, involves an *opening* (upward in this plot instead of downward). That exceptional subject, corresponding to the lowest open circle in the left panel of Figure 21, is subject 10. Her detailed Procrustes decagonal representation, the last panel in the second row of Figure 8, suggests a failure of her depressor labii inferius muscle — the lower lip simply does not lower as far as every other subject’s does. A different projection, in Figure 19, shows that this subject’s principal direction of open-lip shape change in shape space is indeed different from everybody else’s.

The perfect separation of these two bundles of smiles in our subspace projection, in turn, suggests a further step, the creation of spatiotemporal shape coordinates (STSC’s) that treat these points of the trajectory as if they were themselves digitized reference points in some geometry that could be converted to their own shape space and explored further there. The STSC’s here are the equivalent of the *two-point* or *Bookstein* coordinates derived from the original geometries in Figure 21. Figure 22 shows the result, again in two frames corresponding to our two experimental conditions. At right we see that once we have standardized for the net displacement of the projected shape over the course of the closed-lip smile, the trajectories that result are quite similar to one another all along their length, just what we might expect from a behavior that is actually expressing the contraction of one principal muscle, the zygomaticus major. By contrast, the corresponding superposition plot for the open-lip smiles, at left in the figure, is wildly heterogeneous.

**Figure 22.**
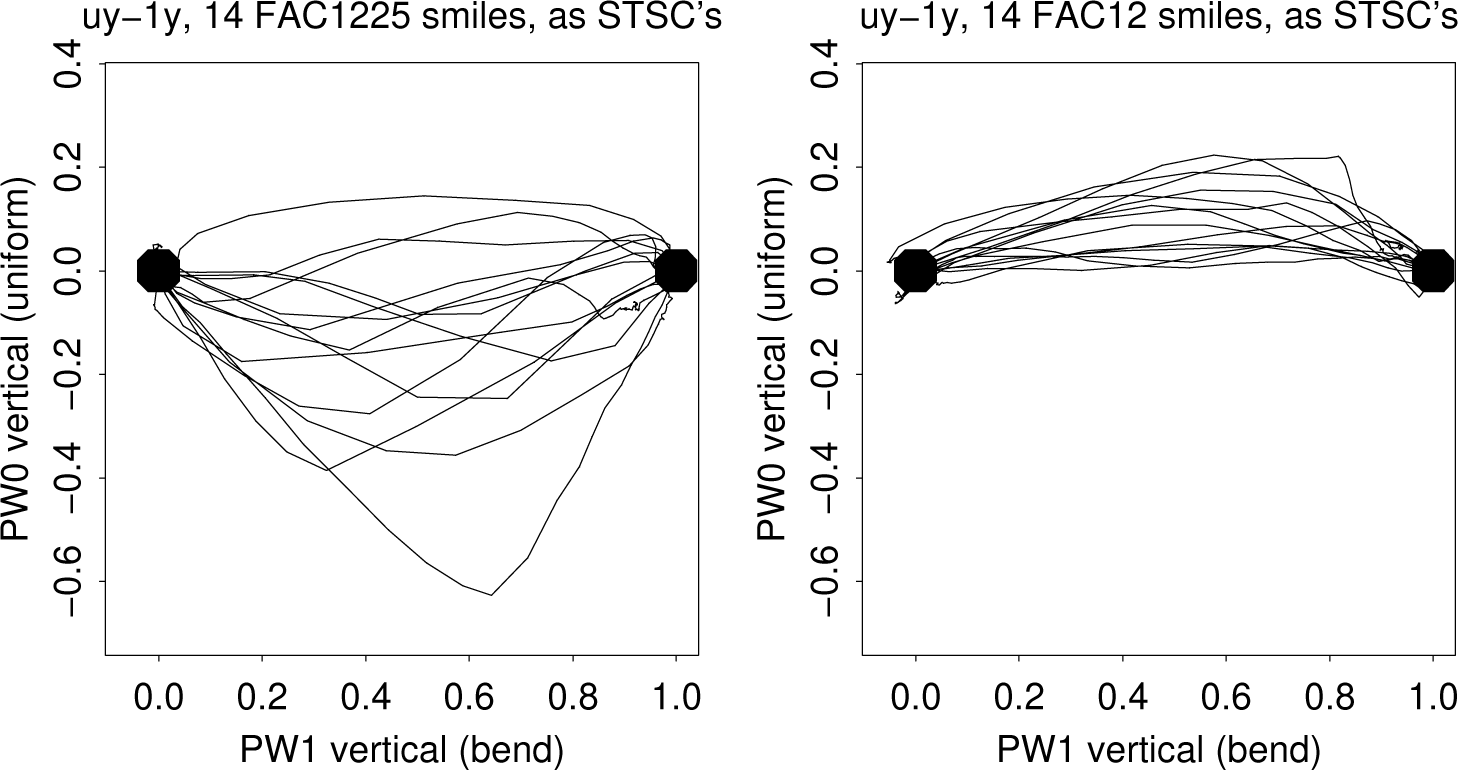
Further standardization of the trajectories in Figure 21 by conversion to spatiotemporal shape coordinates (STSC’s). The closed-lip smiles, at left, seem to vary by only a single additional parameter, the curvature of this trajectory (or, equivalently, the height of the midpoint), corresponding, we think, to nonlinearity in the mechanical consequences of the action of the zygomaticus muscle. The open-lip smiles, at right, vary far more widely. In a larger sample of these smiles we might well be able to uncover interpretable subtypes.

The horizontal coordinate in Figure 22 corresponds to the parameter of “projected motion along the geodesic” explored in Faraway and Trotman (2011). But the vertical coordinate here, which clearly contains valuable information (as its mean is always greater than zero for one group of smiles and always less than zero in the other group), goes unrecognized in their work, in which the analogous projection is interpreted as unstructured variation instead (p. 748). Of course this discrepancy might be due to differences in the data base in addition to differences in methodology (the preservation of all the dimensions of Procrustes shape space, not the reduction to just the two of largest scale as we have done here).

Our principal findings are explicit in the combination of Figure 21 and Figure 22. Within this carefully engineered two-dimensional subspace of the ordinary Procrustes shape space for these 28 series of dynamically deforming decagons, the separation of the two kinds of smiles achieved, the closed-lip and the open-lip, is perfect (Figure 21). Furthermore, while the closed-lip smiles all seem to be quite similar in the hint of a one-muscle biomechanical trajectory, those for the open-lip smile show greatly increased variability (more clearly in Figure 22 owing to the lessened degree of graphical entanglement) owing to the combination of two muscle actions, contraction of the zygomaticus along with relaxation of the orbicularis, that usually appear to be asynchronous in timing as well as highly variable in relative magnitude.

Statistical confirmation of these claims is straightforward. The null hypothesis of no separation of the two clusters of points along the vertical coordinate in Figure 21 shows a *p*-value of 2 × 10^−6^ by t-test and 5 × 10^−8^ by Wilcoxon test (an order statistic). Over in Figure 22, a test for mean difference in the height of these STSC trajectories, now further truncated to the middle 80% of their ranges here, yields a tail-probability on the null of 0.00051 for equality of path mean heights (by t-test) and 0.00027 (by the standard F-test) for equality of variances of those path means between the two classes of smiles. Clearly, in either representation the claim of a group difference meets the “interocular traumatic test” (Bookstein, 2014:xxvii) — the pattern hits you between the eyes.

#### … which in turn align perfectly with a-priori anatomical knowledge

All the preceding considerations can be summarized in Figure 23. The grids in the top row depict what has emerged as factor 1 of our anatomical explanations. It corresponds to a steady extension in direction 1*y* of the shape space, which is a bending along the horizontal axis. This is, of course, a consequence of the action of the zygomaticus major muscle, which acts (see Figure 3) roughly along this direction on the solid skull. In the second row of the figure is a sketch of the action of factor 2, the opening of the lips, corresponding to the relaxing of the orbicularis oris muscle, whenever it happens to be triggered later in the smile trajectory. This delay of triggering is expressed in the figure as the identity of the first two frames in this row — the orbicularis muscle remains quiescent for an interval after the initiation of zygomaticus action, an interval that, were this a larger sample of smiles, could itself be estimated statistically.

**Figure 23.**
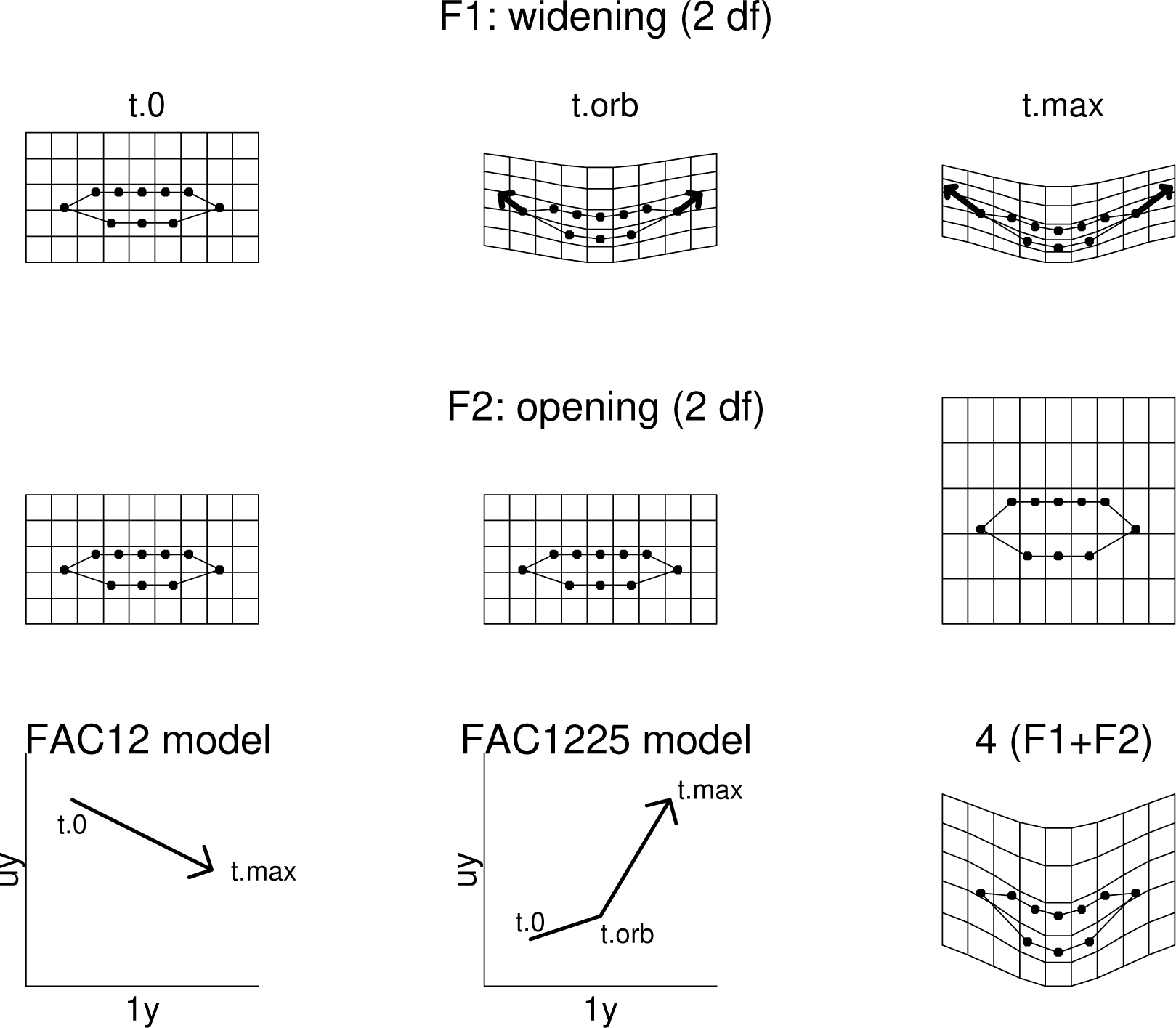
Schematic of our interpretation of the 1*y*–*uy* projections. Top row: the closed-lip smile, corresponding to partial warp score 1*y* and also to the action of the zygomaticus major muscle. Heavy arrows: graphically shifted vectors showing the displacements responsible for this shape change. The apparent vertical compression is a Procrustes artifact of this increase in horizontal diameter. Second row: the lip-opening component of the open-lip smile, corresponding to the action of the orbicularis oris muscle. The columns in these two rows correspond to the three times of interest in the analysis (start, change point, completion of the smile in question). Bottom row, left and center, the corresponding schematics in the space of Figure 21; right, net effect of the FAC1225 (sum of the two grids above it, exaggerated by a factor of 4). *t.*0: time smile dynamics begins. *t.orb*: time of onset of orbicularis relaxation. *t.max*: time of completion of smile dynamics.

The third row of the figure combines these two postulated spatiotemporal patterns as a pair of prototype trajectory models in their common projected (*uy,* 1*y*) subspace. The FAC12 model, drawn here as a straight chord, is actually typically curved in the figure, perhaps owing to actions of other muscles than the zygomaticus in the lip corner displacement; we do not model that curvature here. That for the lower right sketch represents the behavior of the five specific sequences in Figure 19 that show a distinct change point where the second muscle action (orbicularis relaxation) clearly begins well after the contraction of zygomaticus. the orbicularis opening and the time of its triggering. In future modeling this time series, too, will gain an additional parameter.

The variability of the FAC1225 smile that we highlighted at right in Figure 21 or Figure 22 can be illustrated less abstractly by careful selection of a pair of individual smiles to contrast. From the Figure 21 scatter of trajectories we select two: the one making the sharpest angle in Figure 19 — this is subject 7 — versus the subject who arrives at the same final change in Figure 21 but happens to get there by a strikingly linear trajectory instead. This is subject 02, whose trajectory in Figure 19 is indeed remarkably close to flat. Figure 24 retains the coordinate system of Figure 22 but shows only this specific pair of trajectories.

**Figure 24.**
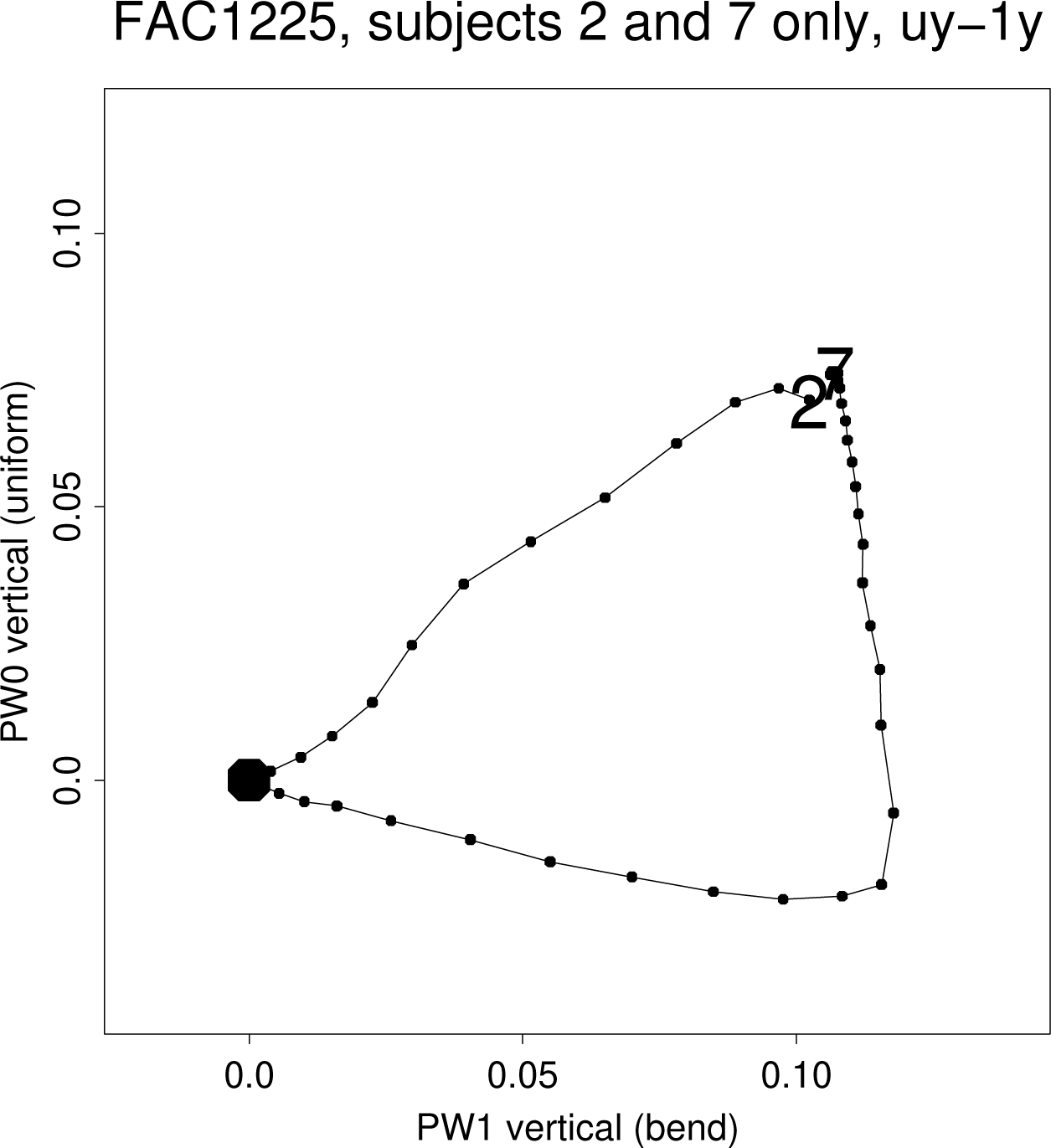
Contrast of the trajectory having the clearest change-point, for subject 7, with a flat (monotone) trajectory, subject 2, arriving at the same net displacement in this projection. Dots correspond to individual resampled frames.

When dynamics are this different we ought to be able to visualize the situation by less complex, more familiar aspects of GMM back in the coordinate system of the original images and their ten-landmark representations. For subject 7, the corresponding three original video frames are spread out in Figure 25 in roughly the geometry of Figure 22. Their images have been augmented by the standard thin-plate spline on all three edges of their triangle: start to middle (left side), middle to final smile (right side), start to finish (top edge). Evidently the change from starting form to change-point is entirely different in biomechanics from the subsequent change to the end of the trajectory. Namely, the first spline extends the corners, as in the rightmost panel of Figure 23, whereas the second spline opens the lips, as in the central panel of that figure. For subject 2, the grids for the two halves of this smile (not shown) appear instead to be virtually identical, corresponding to the straightness of the trajectory in Figure 24.

**Figure 25.**
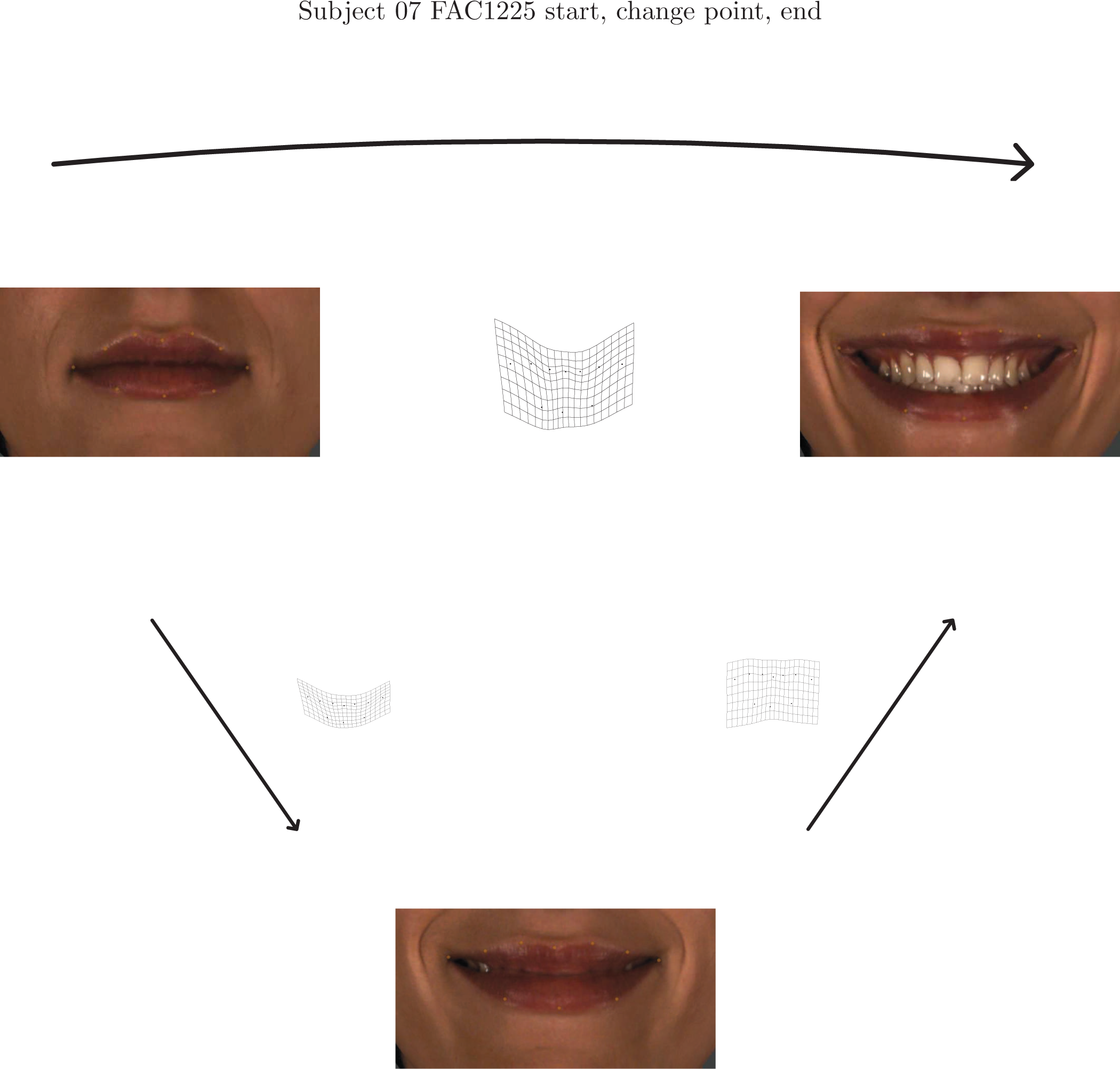
The trajectory for subject 07 from Figure 24, reduced to just three key frames positioned roughly as they would have been in Figure 22, along with the grids for the initial motion segment (left), the final motion segment (right), and the net smile change overall (top row). The segmental grids are obviously different. All grids have been exaggerated twofold for legibility.

#### Recapitulation

We close this survey of results by returning to Figures 3 and 4, embodying the original GMM approach to subject 07’s smiles, in the context of this newly anatomically grounded understanding. In Figure 3, the zeroth and first partial warps (left two columns) apparently represent the actions of two different muscles, orbicularis oris and zygomaticus major, respectively. These two components dominate the dynamics of either smile, the open-lip or the closed-lip. In the closed-lip smile, they are synchronous; but in the open-lip smile, the orbicularis component begins its action long after the zygomaticus component has begun its action. Generalized over all fourteen subjects, Figure 21, the synchronization of the dominant muscle pair in the closed-lip smile is universal, but the dissynchrony for the open-lip smile is heterogeneous. For this specific subject, the transformation corresponding to the first segment of the smile is one of zygomaticus contraction only (Figure 23, upper row; Figure 25, grid at left), while the second segment is mainly a relaxation of orbicularis (Figure 23, middle row; Figure 25, right-hand grid). For many of our other experimental subjects this second phase appears to be a combination of both factors, tht is to say, both muscle actions, not just one at a time.

Thus while the partial-warp plots in the left column of Figure 4 are coherent with the qualitative understanding of this dynamics supplied beforehand by medical anatomy, the plots of relative warps in the right-hand column, which embody the standard GMM approach (Dryden and Mardia, 2016), are not. For the open-lip smile, they represent an anatomically arbitrary rotation to a pose of greatest explained Procrustes variance, which is not a biologically meaningful quantity (Bookstein, 2017a); for the closed-lip smile, the rotation conceals the synchronization of the two muscle actions inside the synthetic descriptor that is their sum, while the second dimension represents an arbitrary selection from the remaining six dimensions of the lower row in Figure 3. Throughout this exemplary little study, to arrive at a proper anatomical understanding of muscle dynamics, with all the associated implications for cognitive and evolutionary neurobiology, requires a deep reconfiguration of the GMM toolkit in order to afford a possibility of identifying statistical factors with anatomical interpretations.

### IV. Discussion

We reported two basic findings. The closed-lip and open-lip (maximal) smiles involve entirely different lip shape changes, and the closed-lip smiles are a fairly well-behaved ensemble of examples over our sample of 14 subjects, whereas the open-lip smile is heterogeneous not only as regards the outcome (the smile actually performed) but also as regards the trajectory of shape changes by which the lips arrived at that ultimate shape — different enough that probably no statistical techniques such as averaging can actually be justified physiologically. This is a purely morphometric description, but we have emphasized throughout that the two dimensions of the plots onto which we have projected our sample in the course of pursuing this summary correspond closely to the known division of labor among the facial muscles responsible for these forms and the known classification of the two types of smile explored in this experiment. This degree of alignment, we assert, accomplishes the bridging of GMM to medical anatomy that we promised in our Introduction. Most of this discussion deals with the import of that unusual extent of transdisciplinary matching. A very brief overview of this analysis is given in Mardia et al. (2018).

#### On the language of a bridge from GMM to anatomy

A latter-day Thompson would appreciate this example in the spirit of the original *OGF* of 1917. Figure 20, the net change, is either one integrated change (the closed-lips subsample) or the sum of two of them to a varying ratio (the open-lips subsample). Each of these meets Thompson’s requirement that it *separately* suggest “forces” (in this case, arising from muscle actions at some distance from the data frame). In identifying these features with anatomical entities we have explicitly chosen to leave the small-scale patterns of change (the rightmost few columns in Figure 3) as obscure to the eye as they would have been in 1917. Over the century, the main desideratum of the biometricians attending to this nexus has been the separation of components of this “integrated change” at their different scales, and our 21st-century GMM praxis is suitable for this purpose should the data afford it. If they are relevant to the health or evolution of the organism, today’s researchers would explore their origin in differences of innervation or mechanics among all the facial muscles, a rhetoric of explanation to which neither Thompson nor the present authors had access.

Most of the founders of GMM were attempting to optimize it as a tool for biological investigations — for the production of graphically intuitive quantitative descriptions of real organismal processes — rather than for estimates of a-priori scalar parameters or the stereotyped “significance levels” associated with elementary null hypotheses of no biological importance. Explanatory purposes like these have implications for the statistician’s choice of a basis for the representation of findings (and their diagrams) in landmark shape space. A redundant basis of shape coordinates of single landmarks separately, while adequate for some tests of null hypotheses, makes no scientific sense in view of the constraints on those sets imposed by the normalizations that are part of the Procrustes algorithm (Bookstein, 2016) and also by the reality of morphological integration (Bookstein, 2015a). The purposes of a biological investigation can be served well only when this basis is rotated to a set of dimensions that can be interpreted as potentially independent, separately meaningful processes whenever the arithmetic justifies such an interpretation.

The partial warps of the data analysis here comprise just such a biologically cogent basis, and when a subsequent factor analysis demonstrates them to align adequately with the contrasts of interest (Figures 15, 16; see Bookstein, 2017a), then that rotation itself becomes justified on biological grounds just as the analogous factor rotations have long been justified in psychometrics on explicitly psychological grounds. It is not the shape coordinates that bear the interpretation of GMM findings, nor even their principal components, but instead the rotations into the separately interpretable dimensions corresponding, ideally, to separable physiological processes (in well-defined physiological experiments like this one), to separable selective processes (in well-defined evolutionary comparative analyses), to separable auxological processes (in well-designed studies of growth), and so forth. In statistical terms, we have demonstrated a fusion of GMM with one concern of classic factor analysis (see Lawley and Maxwell, 1963), the representation of relationships among the variables of a suite, and extended it to the arithmetic of varimax rotation. In the language of biometrics, the same example may be described as retrieving an appropriately contrasting pair of quantitative pattern descriptions from an experimental design aimed at precisely that outcome.

#### Limitations

The limitations of this study are mainly those common across the genre of methodological experiments in general, including all of the examples put forward in the founding papers of GMM as it exists today. Our sample size is small, so we cannot even explore sexual dimorphism of these smiles. (But once we arrive at Figure 21, the count of meaningful variables, namely, 2, falls far short of our count of 14 cases, permitting a powerful inference indeed, namely, a perfect separation over the two experimental conditions being contrasted.) We have no data on actual muscle origins or insertions, let alone electromyographic records of the tempo of their various contractions; such data would be far too invasive for an exploratory study such as this, especially in view of the fact that all of our subjects are physiologically normal. Likewise we have omitted any investigation of the information in the last six partial warps (e.g., the rightmost six columns in Figure 3). Figure 17 shows that the magnitude of these fugitive terms is occasionally substantial, though not reliably so. Surely these account for many of the features that allow us to recognize particular smiles on particular faces, such as actors’. The BE–PwV plots, Figures 13 and 14, tell us that our two selected dimensions *uy* and 1*y* of shape change exhaust at least 80% of the shape variation displayed in 27 of these 28 smiles, but that is not to say that the residual is without psychological significance.

Keep in mind, furthermore, what the overarching goal of this investigation actually *was*: to build a bridge between GMM and a-priori medical anatomy in a suitable first context, so as to illustrate Thompson’s mantra of “the origins of form in force” in a physiologically realistic way. The data we analyzed involved two systems of forces and a corresponding collection of dynamic sequences of forms; and we showed that the comparative statistics of the pair of systems indeed matches the change of forces built into the measurement design. For both of these smiles, “the form of the entire structure under investigation [is] found to vary in a more or less uniform manner,” as Thompson put it. We thereby learn *both* that today’s GMM of systems varying “after the fashion of an approximately homogeneous body” is tuned to analysis of integrated partial warps — cf. Figures 11 and 12 — and that inhomogeneity of *samples* (a concern that entered morphological statistics, alas, only well after the date of Thompson’s first edition) distinguishes well between the settings of systematics versus physiology.

In this regard our demonstration here seems to have gone beyond the styles of quantifying facial expressions in the current literature. Analysis of trajectories in shape space (e.g., Adams and Collyer, 2009), though suggested by a reviewer of an earlier draft, is inappropriate for this context, as it fails to distinguish among uniform features, large-scale features, and small-scale features, and so is obviously ill-suited for studies of soft-tissue deformation; perhaps it might be useful in studies of gait or other aspects of articulated linkages. The common core of all the GMM investigations of smiles reviewed in Section I.2 above is evidently misspecified in ignoring the overwhelming evidence of integration among these decagons of shape coordinates. To demonstrate their association with anatomical knowledge requires a method that acknowledges the anatomical knowledge of this integration as conveyed a-priori in atlases as well as in the FAC system itself, which classifies expressions by their muscular origins. In our view the method we have demonstrated here — the mapping of anatomical prior knowledge against a factor structure revealed by GMM’s extant scheme of partial warps — is the only current candidate for erecting this bridge.

Then the characteristics of the present example are not so much limitations of the inferences in this paper as possible limitations of the generalizability of the method it embodies. This is, indeed, a new way of pursuing the multivariate statistical analysis of a space of Procrustes shape coordinates. It gets the right answer (a pair of dimensions matching the a-priori understanding of the experiment) here, in this setting that comes so close to the original spirit Thompson set down a century ago. So it might well apply to other settings that meet the same requirements — other studies of form change that originate in actual forces, realized in experimental designs that effectively control those forces. And when heterogeneity is found to be irreducible by these approaches, as we found for the relative weight and timing of the pair of muscle actions in the FAC1225 condition, the more intensive measurements needed to quantify *these* concerns, and perhaps their abnormality in the presence of abnormalities of form (such as the smiles of persons with repaired clefts), become much easier to justify.

The geometrical directions associated with the Procrustes displacement of lip commissures in the closed-lip smile obviously do not correspond to the real lines of action of these muscles, as often they point inward in Figure 10 whereas the real muscle action is outward. Discerning actual lines of muscular action requires information that has been obliterated by the Procrustes maneuver, as these lines of action would need to be specified by structures at some distance from the decagon of fiducial points on which our calculations here rely. Now that we suspect that the trajectory of the FAC12 smile is representable fairly closely as the homogeneous action of one single muscle, we can ignore the evidence of integration in the slopes of the little line segments fitted to the bending energy – partial warp variance plots in Figure 13. Such a further investigation would require analysis by the tools of biomechanics, not shape geometry: for instance, representations of the layers of facial epidermis and dermis and their bulk material properties such as compressibility or hysteresis. The parameters of such representations are retrievable only from the tools currently known as “dense phenotyping,” not from a mere handful of artistically familiar loci along a single high-contrast anatomical curve. Similarly, while the orbicularis oris muscle is actually a closed circuit (four interwoven quadrants of fibers) around the same outline we have been drawing as a decagon, there is no evidence to that effect in the analysis presented here. That its action in the setting of the open-lip smile is a relaxation rather than a contraction is not retrievable from these data, nor that that relaxation is azimuthal all the way around the outline. What we *can* see, from Figure 14, is that typically this action is not integrated around the circuit. It is safe to treat the residual displacements of these 10 landmarks as independent once we have partialled out the primary factors represented in the plane of *uy* and 1*y*.

In an appropriate GMM analysis of any complex organismal phenomenon, the process of discovery is not sequestered to any single post-arithmetical stage of interpretation. Instead it drives the actual *production* of our dual system of diagrams here for patterns in the closed-lip and then the open-lip smile conditions. We needed to continually inject prior physiological knowledge, even if qualitative, in order to steer the series of statistical maneuvers. In Mardia’s famous model of the elephant (e.g. Dryden and Kent, eds., 2015, cover art), those observers of an animal in a zoological park are talking to one another, so that each one’s questions depend on the reports of those who investigated earlier.

#### Biotheoretical aspects of the bridge

Such interplay between arithmetic and understanding is embedded deeply within the foundations of comparative anatomy. As Elsasser (1975) trenchantly notes, from his vantage point within the neighboring field of theoretical biology,

> In physical science one deals invariably with definite abstract constructs, that is, sets of symbols, most commonly mathematical, in terms of which observed regularities can be represented. … Certain complications arise in modern atomic and molecular science whenever concepts of probability enter. This indicates that one no longer deals with individual objects but only with classes of these. Once the transition from objects to class is made, the abstract scheme of the description that uses probabilities will, in terms of classes, become as precise as the direct quantitative description of single objects in classical physics.

> But there is one exception, the description of objects that are of radically inhomogeneous structure and dynamics, which we identify with living objects. Here a property of the description that can otherwise be ignored will be of the essence: it is that for a somewhat complicated object the number of possible microscopic patterns compatible with a given set of known data (class parameters) becomes immense. … The description of a living object, therefore, implies an *abstract selection.* It is the selection of an immensely small subset from an immense number of possible descriptions. … One goes from theoretical physics into theoretical biology not by the method of superadding new, formal constructs (an approach usually described as vitalism) but by omitting an overwhelming fraction of an immense number of available descriptions. [page 203; emphasis in original]

GMM is part of Elsasser’s domain here — it is a tool for description of “objects that are of radically inhomogeneous structure and dynamics.” We needed the models of scaling dimension, Figure 18, to bifurcate our attention between an isotropic component, promptly to be discarded just as a type of error, and a signal space, which, conveniently enough, is realized within the largest-scale four-dimensional subspace of the shape space according to the spectrum of TPS bending energy that is shared across all of today’s best GMM applications. We further focused on the plane within that subspace representing only the bilaterally symmetric processes, sequestering the asymmetric component of equal dimensionality (two: *ux* and 1*x*) as part of the “noise” that this study was not designed to analyze.

Needless to say, the “bending energy” of the splines here has no relationship to the actual energetics of the facial muscles that actually produce these changes. Indeed, the open-lip smile involves a relaxation, that is to say, a physiological energy that is actually *negative* with respect to the the neutral “resting” expression. Such niceties of the relation between statistical arithmetic and actual biomechanics are, again, inaccessible from the study laid out here. It is not only the limits of the GMM toolkit that block this deeper analysis, but the incompatibility between the GMM and the anatomical notions of reductionism. In GMM, increasing the range of available scales adds detail to the derivatives that are our currency of signal strength. But in the actual anatomical sciences, this same process of “drilling down” adds entirely new levels for the representation of process (tissues rather than empty picture spaces). GMM and anatomy have entirely incompatible notions of “explanation,” and those of anatomy must be granted priority (Bookstein, 2017a, 2018).

The grid for PW1y that Figure 20 or Figure 23 displays is a substantial smoothing of the actual FAC12 shape trajectory of the data decagon, one that suppresses many details. Although it highlights the responsible facial muscles correctly as they relate to the experimental design (contrast between open-lip and closed-lip smiles), it is unrealistically symmetrical, not only left-to-right but also upper-lip-to-lower, owing to the analogous symmetries of the landmark configuration per se in Figure 5. Yet that top-to-bottom symmetry is actually broken in these data; this is clear in some of the subordinate principal warps in Figure 3, particularly PW4 and PW7. On the other hand, the stasis of Procrustes position for semilandmarks 2, 6, 8, and 10 in Figure 9 suggests that the near-linearity of PW1y along the corresponding polylines 1-4, 4–7, 7–9, 9–1 of the lip outline might actually be valid. In this linearity the spline model here disagrees with another current tool, the analogous parabolic model of Bookstein (2024). (That model, a refinement of one originally suggested by Sneath in 1967, likewise was intended explicitly to supersede Thompson’s subjective “circles” and “hyperbolas” reproduced in our Figure 1.) Both of these later approaches go well beyond Thompson in foregrounding a whole new *class* of tool, a finite *spectrum* of descriptors that increase in detail but generally decrease, one hopes, in signal amplitude. In applications to actual physiological processes, such as the muscular contractions resulting in these smiles, one searches first for a match, however rough, between the a-priori understanding of muscular actions at large scale and the corresponding large-scale geometric responses. Any matches detected are subject to modification by smaller-scale details of geometry paired with smaller-scale aspects of anatomy (in this instance, other facial muscles whose effects are too fugitive to be assessed by a sample of fourteen subjects only, or perhaps aspects of tissue compressibility or anchorage specific to particular arcs of the lips inside the vermilion border traced here).

#### Future directions, and final thoughts

We envision two extensions of this work that will require small modifications of the methodology just reviewed. One such extension, the original context for which this preparatory work was envisioned, is the study of the effects of cleft palate repair surgery. In this medical context, surgeries that interfere with the actions aligned with the factors here may well have effects on their geometric expression, not only the histology of their scars. Clefting entails a catastrophic change in the action of the orbicularis muscle, and we should be able to see that reflected in major alterations of the factor structure we imputed for these data. Because clefting is typically unilateral, however, we would expect analysis to benefit from a rotation to the basis of symmetric and asymmetric components of bilateral structures like this one. Such a rotation is already standard in the GMM literature, but attempts at the geometrization of a subjective rating of unattractiveness may benefit if, for example, they prohibit the rotation of asymmetric structures with respect to laboratory coordinates that lies at the basis of the standard Procrustes approach.

Additionally, for analyses of attractiveness and other psychological correlates of the anatomical dynamics here, there may be information in the actual scaling of a smile beyond the shape patterns analyzed in this paper. The extension of all these results to *form space,* the augmentation of a Procrustes shape analysis by one additional dimension that restores the original geometric scale by its logarithm, is likewise a straightforward application of current techniques.

Representation of smiles is also possible in a system registered on the immobile structures of the upper skull (orbits, forehead). Studies exploiting that registration typically assess net displacements of lip landmarks point by point (e.g., Sidequersky et al., 2016); but this vitiates all the statistical benefits of anatomical integration borne in the dominance of PW0 and PW1 in our analysis. Specialized analyses like these are valuable for quantifying the reliability of low-level image processing protocols (e.g., Ju et al. 2016), but to our knowledge they lack the discriminatory power of the low-dimensional shape space projections such as that in Figure 21 whenever biological hypotheses about distinctions among organismal conditions are to be unearthed. We invoked neither of these extensions here: not the study of asymmetry (e.g., Mardia et al., 2000), because the smiles in our sample are overwhelmingly symmetric; not the extensions to the unscaled coordinate systems, because nothing in the literature of such analyses matches the power of our basic 1*y*-*uy* finding here.

Our basic finding was a match between the statistics of the representation of two smiles and prior anatomical knowledge of the distinct muscular origins of those two stereotyped sequences. We showed that an experimental study of two different smile conditions, one produced as an action primarily of one muscle and the other the action of two, can reproduce the physiology of the comparison by a pair of statistical summaries of the corresponding partial warp scores. While the standard toolkit of geometric morphometrics is not equipped to analyze a data set of real muscle actions in a biomechanically sensible fashion, nevertheless these novel extensions borrowed variously from evolutionary biology (the scaling dimension) or psychometrics (factor analysis) made it possible to build a bridge to an appropriate understanding. Biological shape per se, after all, is just a measurement, not an embodiment of any theory. As Elsasser implied, the purpose of any quantification in a biological science is to *limit* the range of possible explanations that we might pursue to those likeliest to convey a scientifically meaningful message. We model biological processes by deleting information, not by accumulating it. Our example is perfectly in keeping with this dogma of Elsasser’s, inasmuch as the summary diagrams in Figure 21 referred only to two partial warp scores out of 16 and only to five parameters out of a full multivariate complement of 136 required to specify the general covariance structure of a vector of 16 variables, the count of dimensions of our Procrustes shape space here.

The way a shape analysis serves to deepen our understanding of organisms requires us to apply every bit of intelligence we have about how arithmetic, properly managed, can sometimes lead to that understanding. Crude Procrustes analyses do not lead to an appropriate understanding of the normal human smile — they should not have been expected to. But in combination with other tools, we have managed to interweave our arithmetic with representations of actual anatomical knowledge in order ultimately to arrive at a valid comparison of grouped results from this small but sturdy experimental design.

We commend this approach — the layering of borrowed tools from a wide range of other sciences atop the simplistic arithmetic of a Procrustes fit — as a good model for quantitative anatomical investigations in general. At least in the course of studies of organismal form and its variations, the intentional subordination of statistical arithmetic to prior qualitative knowledge (Bookstein, 2015c) usually leads to more profound scientific progress than any mastery of statistical methodology per se would have done. In other words, the representation of *un*certainty in statistics needs to be based not on probability theory but on the actual *certainties* of scientific knowledge as it actually exists prior to the launching of any systematic quantifications. Our demonstration that in at least one setting we have recovered prior qualitative anatomical knowledge from multivariate statistical manipulations thus may serve as a prototype for the extension of GMM into the anatomical sciences in a variety of other comparative contexts: but only if anatomical knowledge is guaranteed the right to dominate the strategies of statistical manipulation, as it has been in our example here.

#### Consent

All subjects gave us their informed consent for the use of their data in this way. This project was approved by the University of Leeds Research Ethics Committee, School of Dentistry, application 240915/BK/179.

## Acknowledgement

We are grateful to three anonymous reviewers for showing us the many places where the arguments of an earlier version had been ambiguous or incomplete. Collection of the data resource explored in this report was supported by the Faculty of Dentistry and the School of Mathematics, University of Leeds. Mr. Cian Lowney was responsible for subject recruitment and image capture.

